# Integrin αVβ6-EGFR crosstalk regulates bidirectional force transmission and controls breast cancer invasion

**DOI:** 10.1101/407908

**Authors:** Joanna R. Thomas, Kate M. Moore, Caroline Sproat, Horacio J. Maldonado-Lorca, Stephanie Mo, Syed Haider, Dean Hammond, Gareth J. Thomas, Ian A. Prior, Pedro R. Cutillas, Louise J. Jones, John F. Marshall, Mark R. Morgan

**Author notes:** Corresponding author: Dr Mark R. Morgan, PhD, Cellular & Molecular Physiology, Institute of Translational Medicine, University of Liverpool, Crown Street, Liverpool, L69 3BX, UK. Tel: [+44](0) / / Twitter:@M_MorganLab Professor John F. Marshall, PhD, Centre for Tumour Biology, Barts Cancer Institute, Queen Mary University of London, John Vane Science Centre, Charterhouse Square, London EC1M 6BQ, UK Tel: [+44](0) 20 7882 3580 /. Denotes equal contribution (Joanna R. Thomas, Kate M. Moore).

## Abstract

The mechanical properties of the extracellular matrix within tumours control multiple cellular functions that drive cancer invasion and metastasis. However, the mechanisms controlling microenvironmental force sensation and transmission, and how these regulate transcriptional reprogramming and invasion, are unclear. Our aim was to understand how mechanical inputs are transmitted bidirectionally and translated into biochemical and transcriptional outputs to drive breast cancer progression. We reveal that adhesion receptor and growth factor receptor crosstalk regulates a bidirectional feedback mechanism co-ordinating force-dependent transcriptional regulation and invasion.

Integrin αVβ6 drives invasion in a range of carcinomas and is a potential therapeutic target. αVβ6 exhibits unique biophysical properties that promote force-generation and increase matrix rigidity. We employed an inter-disciplinary approach incorporating proteomics, biophysical techniques and multi-modal live-cell imaging to dissect the role of αVβ6-EGFR crosstalk on transmission of mechanical signals bidirectionally between the extracellular matrix and nucleus.

We show that αVβ6 expression correlates with poor prognosis in triple-negative breast cancer (TNBC) and drives invasion of TNBC cells. Moreover, our data show that a complex regulatory mechanism exists involving crosstalk between αVβ6 integrin and EGFR that impacts matrix stiffness, force transmission to the nucleus, transcriptional reprogramming and microenvironment rigidity. αVβ6 engagement triggers EGFR & MAPK signalling and αVβ6-EGFR crosstalk regulates mutual receptor trafficking mechanisms. Consequently, EGF stimulation suppresses αVβ6-mediated force-application on the matrix and nuclear shuttling of force-dependent transcriptional co-activators YAP/TAZ. Finally, we show that crosstalk between αVβ6 & EGFR regulates TNBC invasion.

We propose a model whereby αVβ6-EGFR crosstalk regulates matrix stiffening, but also the transmission of extracellular forces into the cell in order to co-ordinate transcriptional reprogramming and invasion. To exploit adhesion receptors and receptor tyrosine kinases therapeutically, it will be essential to understand the integration of their signalling functions and how crosstalk mechanisms influence invasion and the response of tumours to molecular therapeutics.

## INTRODUCTION

Dynamic interactions between breast cancer cells and the local tumour microenvironment directly contribute to disease progression, prognosis and chemoresistance^1-5^. The mechanical properties of the extracellular matrix (ECM) within breast tumours control multiple cellular functions to drive cancer invasion and metastasis; stiff ECM promotes invasion and correlates with poor patient survival. The extracellular micro-environment of tumours is characteristically stiffer than normal tissue, as a result of ECM deposition, remodelling and stromal cell contractility. It is thought that a self-amplifying circuitry exists, linking tissue stiffness, ECM resistance and cell-mediated contractility. These positive-feedback mechanisms impact tumour cell invasion, transcriptional reprogramming, metastasis and patient survival^1, 2^. However, the mechanisms controlling microenvironmental force sensation and transmission, and how these regulate tumour cell function are unclear.

Integrins are transmembrane adhesion receptors that relay mechanical signals bidirectionally across the membrane, between ECM and the contractile cytoskeletal machinery. Thus, integrins enable cells to sense the mechanical properties of matrix and to exert forces on ECM to control tissue rigidity, cell migration and cell invasion^6^. Integrin-associated complexes (IACs) also function as signalling platforms to spatially and temporally control the propagation of membrane-distal signals. Thus, regulation of cell-matrix interactions and dynamics, controls mechanotransduction, cell migration, microenvironment remodelling and global cell fate decisions^6^. Via these mechanisms, integrins mediate cell invasion, adhesion, proliferation, survival and differentiation, and, consequently, dysregulated integrin signalling or expression contributes directly to cancer development and metastasis^7^.

The pro-invasive integrin αVβ6 is upregulated in a range of carcinomas, from normally very low levels, and overexpression is associated with poor survival in several types of cancer including breast, colon, cervix and non-small cell lung cancer^8-11^. Expression of αVβ6 integrin is an independent predictor of breast cancer survival and metastasis^12, 13^ and drives invasion in a range of different cancers^14^. Thus, αVβ6 is a prognostic indicator/biomarker and potential therapeutic target^8, 13, 14^. Indeed, therapeutic targeting of αVβ6 with inhibitory antibodies has produced promising results in breast cancer models in vivo. Relative to other integrins, αVβ6 exhibits distinct biophysical properties that promote force-generation, rigidity sensing and have the capacity to increase matrix rigidity^15^, suggesting that αVβ6 could play a key role in sensing and regulating extracellular mechanical forces in breast cancer.

Integrin αVβ6 expression is restricted to epithelial cells and is usually only detected on cells undergoing tissue remodelling processes such as wound healing and cancer^16^, suggesting that targeting of this integrin would be cancer-cell specific and reduce potential adverse effects. Our recent *in vivo* studies suggest that therapeutically targeting αVβ6 integrin may represent a novel and effective strategy to treat breast cancer^8,^

Triple negative breast cancer (TNBC) is a highly aggressive tumour sub-type characterised by the lack of expression of oestrogen receptor (ER), progesterone receptor (PR) and no overexpression of human epidermal growth factor receptor 2 (HER2). TNBC accounts for over 10-15% breast cancer deaths annually in the USA^17^. There are limited treatment options available, and most treatments result in deleterious side effects due to their non-cancer-specific activity. Current treatment for TNBC is conventional cytotoxic chemotherapy, as effective targeted therapies are unavailable (8). Hence, there is an unmet clinical need, to develop novel molecular-specific strategies for treating TNBC.

However, in order to design molecularly-targeted therapeutic strategies, it is important to understand the signalling mechanisms that regulate, and are regulated by, the target molecules. This is particularly important when targeting integrins, as co-regulatory mechanisms exist between integrins and receptor tyrosine kinases (RTK) (e.g. growth factor receptors), which spatially and temporally co-ordinates the functions of both receptor families. Consequently, integrin-RTK crosstalk mechanisms modulate downstream processes such as migration, proliferation and apoptosis and contribute progression or therapeutic tractability of disease^18, 19^

We investigated whether αVβ6 integrin regulates TNBC progression and survival. We report an association between high expression of αVβ6 and poor survival in both TNBC and ER-negative breast cancer. To further understand how αVβ6 contributes to TNBC development, we employed two unbiased approaches to dissect αVβ6-dependent signalling mechanisms. First, we performed proteomic analysis on ligand-engaged αVβ6 IACs, to define the signalling networks recruited to sites of αVβ6-ECM interaction. Second, we used a phospho-proteomic strategy to identify kinase activation pathways that were activated following ligand-induced endocytosis of αVβ6. These approaches provided a global view of αVβ6-mediated signalling and, independently, identified epidermal growth factor receptor (EGFR) signalling as a key regulatory pathway associated with αVβ6 function.

Follow up experiments, based on further bioinformatic interrogation of the datasets, showed that αVβ6-EGFR crosstalk regulates reciprocal receptor trafficking mechanisms to regulate receptor bioavailability at the membrane. Consequently, EGF stimulation inhibited the ability of αVβ6 to apply mechanical forces on the ECM. We further demonstrate that inhibition of αVβ6 suppressed actomyosin-dependent contractility, RhoA activity and nuclear shuttling of the mechano-sensitive transcriptional co-activator YAP. Interestingly, in MDA-MB-468 TNBC cells, EGF stimulation also inhibited nuclear translocation of YAP; recapitulated the effect of αVβ6 inhibition. Finally, we demonstrate that there is an association between αVβ6 and EGFR expression in breast cancer and that αVβ6 and EGFR co-operate to drive TNBC invasion.

## RESULTS

### Integrin αVβ6 expression correlates with poor prognosis in TNBC

Tissue microarrays (TMAs) from two separate cohorts (London and Nottingham) totalling over 2000 women with breast cancer were stained for β6 expression and investigated for age and disease subtype. The clinicopathological parameters were published previously^8^. We noted a strong association between high αVβ6 expression and poor survival in all women in the London cohort (Figure 1A-B). However, interestingly, in the Nottingham cohort, the association between reduced overall survival (OS) and high αVβ6 expression was determined to be only significant in younger women (≤55 years, HR=2·26, 95% CI=1·46-3·5, *P<*0·001, Figure 1D), rather than those over 55 years of age. Breast cancer in young women tends to be more aggressive with higher proportions of patients with high-grade and later stage (III/IV) tumours with lower oestrogen receptor (ER)-positivity and overexpression of HER2 than their older counterparts^20-22^. Younger women diagnosed with breast cancer often have a poorer prognosis than older women with the disease, so we focused our further analyses on patients ≤55 years (see Figures S1A-F for analysis of additional patient groups).

**Figure 1:**
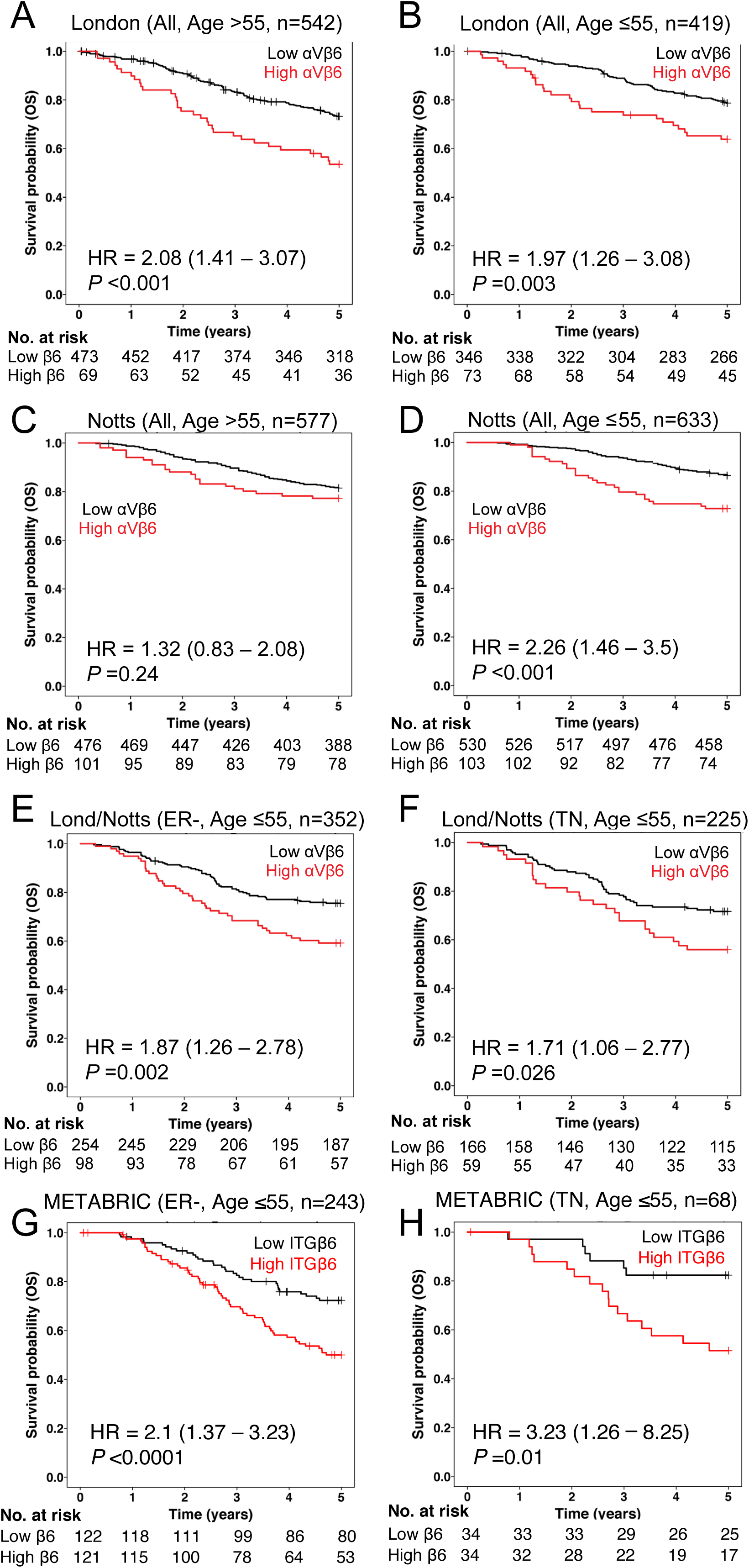
Integrin αVβ6 expression correlates with poor prognosis in ER-negative and triple-negative breast cancer. Kaplan-Meier curves by integrin αVβ6 expression status. Tick marks indicate patients who were censored. All *P* values refer to log-rank tests. Cancerous breast cancer tissue sections were immunohistochemically stained for integrin αVβ6 using 6.2G2 antibody (Biogen Idec). Overall survival in 2 cohorts of breast cancer patients from London (**A/B**) and Nottingham (**C/D**) by integrin αVβ6 status (high αVβ6 expression: red; low αVβ6 expression: black). Overall survival of these cohorts was subdivided into patients >55 years of age (**A** & **C**) and ≤55 years of age (**B** & **D**, termed ‘younger’ patients). (**E**) Overall survival of ER-negative patients (ER-) ≤55 years of age. ER-patients ≤55 years with high αVβ6 expression on their tumours had poorer OS compared to patients with low αVβ6 expression (HR=1·87, 95%CI=1·26-2·78, *P*=0·002). (**F**) Overall survival of triple negative (TN) breast cancer patients from the London and Nottingham patient cohorts ≤55 years of age by integrin αVβ6 status. TN breast cancer patients ≤55 years with high αVβ6 expression showed worse outcome compared to patients with low αVβ6 expression on their tumors (HR=1·71, 95%CI 1·06-2·77, *P*=0·026). Please also see Supplementary Figures 1. **G/H)** Overall survival in METABRIC by ITGB6 gene status (high expression in red, low in black). Overall survival of ER-negative patients (ER-) ≤55 years of age. The *P-*value for ER-patients under 55 years with high ITGB6 gene expression versus low is <0·0001. Overall survival of triple negative (TN) patients ≤55 years of age. The *P-*value for TN patients under 55 years with high ITGB6 status versus low is 0·01

There was a strong association between high αVβ6 expression and poor survival in younger women (≤55 years) in both ER-negative (Figure 1E, HR=1·87, 95%CI=1·26-2·78, *P*=0·002), and TNBC subgroups (Figure 1F, HR=1·71, 95% CI=1·06-2·77, *P*=0·026). This association also was shown at the transcriptional level performing the same analysis on the METABRIC Breast cancer expression database (>2000 cases (24)), where young women with high ITGB6 (integrin β6 subunit) gene expression had significantly reduced survival in the ER-negative (Figure 1G, HR=2·1, 95% CI=1·37-3·23, *P<*0·0001) and TNBC (Figure 1H, HR=3·23, 95% CI=1·26-8·25, *P*=0·01) subgroups. We also observed a significant association between high αVβ6 expression and distant metastasis and OS in the entire Nottingham cohort (Figure S1A, HR=1·38, 95% CI=1·01-1·9, *P*=0·044), Moreover, this association was found to be significant in the ER-negative population of younger women (Figure S1B, HR=1·9, 95% CI=1·1-3·3, *P*=0·019), with a trend towards clinical significance in younger women with TNBC (Figure S1C, HR=1·43, 95% CI=0·72-2·83, *P*=0·301). Together, these data suggest that αVβ6 correlates with poor patient prognosis in both ER-negative and TNBC subgroups.

### Proteomic analysis of αVβ6 integrin-associated complexes

Having established that the pro-invasive integrin αVβ6 is a poor prognostic indicator in ER-negative and TNBC, we sought to delineate αVβ6 signalling mechanisms in TNBC cells. Therefore, we employed IAC isolation techniques and quantitative proteomic analysis to define the signalling networks recruited to αVβ6-dependent IACs.

Three ECM ligands were used for pairwise comparisons to elucidate proteins selectively recruited to αVβ6-mediated IACs. IACs were isolated from the triple-negative breast cancer cell line MDA-MB-468 plated on latency-associated peptide (LAP), fibronectin (FN) or Collagen-I. LAP is an αVβ6-selective ligand, FN can engage multiple integrins including, but not solely, αVβ6, and Collagen-I was used as a non-αVβ6-binding negative control. Prior to proteomic analysis, isolated IAC samples were validated by immunoblotting. Specificity of enrichment was confirmed by the presence of β6 integrin, vinculin and talin, and the absence of non-adhesion-specific proteins (i.e. GAPDH, HSP90 and BAK) in the enriched IACs.

Validated IAC enrichments were subjected to analysis by mass spectrometry to enable global unbiased analysis of the protein network recruited to ligand-engaged αVβ6 integrin. To ensure the quality of the proteomic data, reproducibility across datasets was assessed (Figure S3A-C), peptide identification cut-offs applied, and an overview of fold-changes of protein detection on different ligands was established (Figure S3D/E). For further details, see Supplementary Results Section: Quality assurance of proteomic datasets.

Meta-analysis of previously published proteomic IAC datasets has led to the definition of the proteomically-determined adhesome; comprising a consensus adhesome and the meta-adhesome^23^. The consensus adhesome comprises 60 proteins that were consistently identified as being recruited to IACs and therefore likely to represent core adhesion machinery. The non-canonical meta-adhesome constitutes 2,352 proteins that are more variably detected in IACs; possibly due to lability, highly dynamic/transient recruitment, low stoiciometry, or cellular and ECM ligand context. Interestingly, when we compared our proteomic IAC datasets with the curated consensus adhesome^23^, our dataset contained all of the high-evidence direct integrin interactors in the networks (α-actinin, talin, tensin, kindlin and filamin), but only a small proportion of other proteins within the consensus adhesome (total coverage of consensus adhesome: 35%, 21/60 proteins). Suggesting that the core structural machinery of IACs is conserved, but that other regulatory proteins are divergent when IACs are formed on different ligands. Similarly, the total coverage of the meta-adhesome by our dataset was 20% (468 proteins). However, 30% of the dataset was not represented in either the consensus or meta-adhesome, suggesting the recruitment of novel proteins to LAP- and collagen-engaged IACs, not identified in previous studies. For further details, see Supplementary Results Section: Comparison of IACs to the literature-curated adhesome.

To characterise the αVβ6-associated adhesome, by identifying compositional changes in IACs formed on different ligands, statistical tests (ANOVA of normalised spectral abundance factor values) were applied to determine statistically significant changes in protein enrichment in IACs on specific substrates (Figure 2C). A protein-protein interaction network was then constructed of statistically significant molecules, that were identified as more than 2-fold enriched on LAP, compared with other substrates (Figure 2D). It is important to note that this PPI network is not very well inter-connected (48% of proteins unconnected), because some important IAC structural proteins will be common to IACs on all substrates and therefore not exhibiting >2-fold enrichment on LAP.

**Figure 2:**
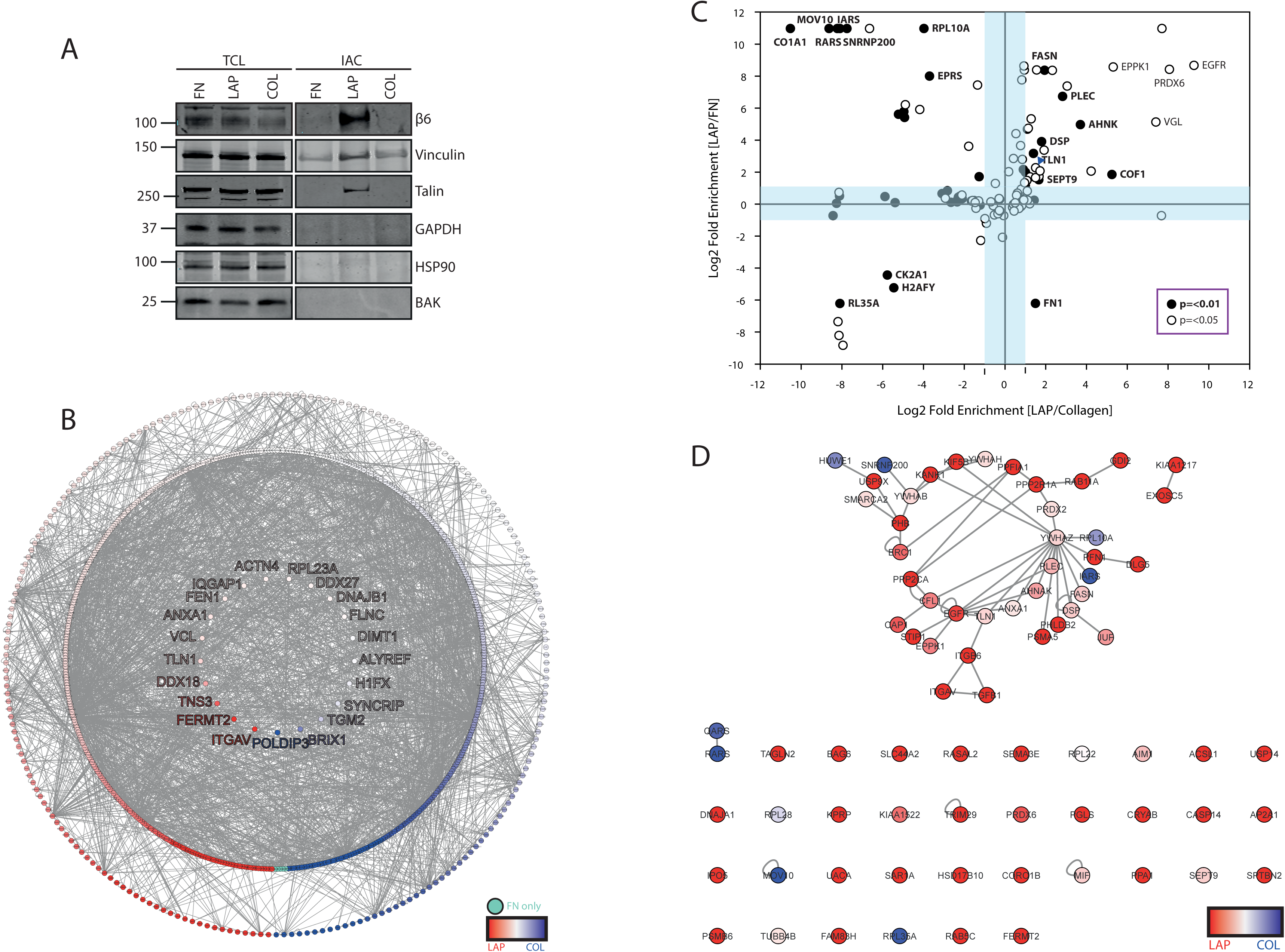
Proteomic analysis of ligand-engaged αVβ6-dependent adhesion complexes. **A)** Validation of integrin-associated complex enrichment. Immunoblotting total cell lysate (TCL) and isolated integrin-associated complexes (IAC) for the adhesion complex components (integrin β6, vinculin & talin), and negative control proteins (GAPDH, HSP90 and BAK from other subcellular compartments).**B)** Representation of consensus- and meta-adhesome dataset coverage.35% of the consensus adhesome (21 proteins), inner ring; 20% of the meta adhesome (468 proteins), middle ring. Proteins in the outer ring were not reported in the consensus or meta adhesome (219 proteins). Total number of proteins = 726. Nodes represent proteins and edges are known interactions. Node colour red to blue gradient = log2 fold enrichment on LAP versus collagen. Green = FN specific proteins. **C)** Statistically significantly different proteins between ligands. Analysis of variance (ANOVA) between LAP, FN and collagen ligands normalized spectral abundance factor (NSAF) values. Statistical analysis was performed in Scaffold (version 4). Statistically significantly different proteins (p=<0.05) were mapped by log_2_ fold enrichment of LAP/ FN and LAP/collagen. Light blue shading corresponds to ≤ two-fold change. Black and white circles correspond to a significance value of p=<0.01 and <0.05, respectively. Gene names of key proteins are indicated in bold or regular text for p=<0.01 and p=<0.05, respectively. Blue arrow head indicates data-point for TLN1. **D)** Interaction network of proteins statistically significantly different between ligands and enriched on LAP.Proteins arranged in a hierarchical ring network. Nodes represent proteins, and edges are known interactions. Node colour red to blue gradient = log_2_ fold enrichment LAP/collagen. Proteins that are enriched on collagen compared to LAP (represented in blue) are positively enriched on LAP compared to FN.

Functional enrichment analysis was performed on proteins significantly, and more than 2-fold, enriched on LAP. Over-representation of gene ontology (GO) term analysis was performed with KEGG (Kyoto Encyclopedia of Genes and Genomes) and Reactome pathway terms. GO term grouping was used to combine related terms into groups. KEGG analysis identified enrichment of proteins associated with ErbB/EGFR signalling, cytokine-cytokine receptor interactions, Hippo signalling pathway, arrhythmogenic right ventricular cardiomyopathy (ARVC), MAPK signalling, fatty acid biosynthesis and endocytosis (Figure 3A/B).

**Figure 3:**
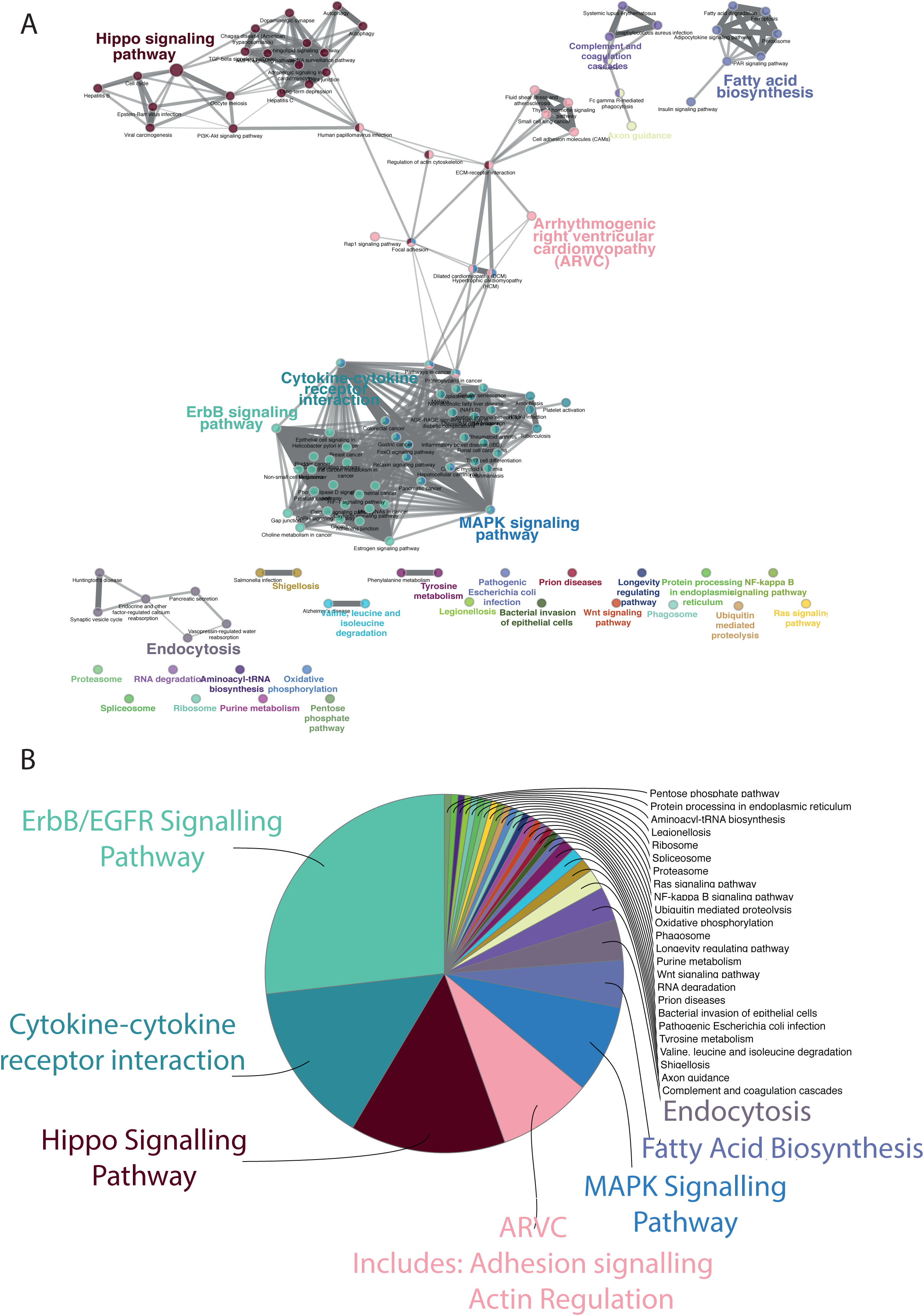
Ontological analysis of αVβ6-dependent adhesion complexes identifies αVβ6-EGFR crosstalk mechanism. **A)** ClueGO KEGG Pathway term hierarchical clustering. Hierarchical layout of KEGG terms identified for proteins enriched in αVβ6-dependent IACs on LAP. Node colour corresponds to grouping, with the lead term in corresponding coloured text. Nodes with split colours belong to multiple groups. Nodes represent individual KEGG pathway terms. **B)** Pie-chart organised by percentage of genes per term. Top 7 lead terms highlighted by colour-coded text. Analysis parameters: minimum 1 gene per cluster, GO term/pathway network connectivity (Kappa score) =1; Statistical test Enrichment/Depletion (Two-sided hypergeometric test), Bonferroni p-value correction; GO term grouping based on kappa score, 50% of genes for group merge, 50% terms for group merge. Leading group term based on % gene/term.

Interestingly, as ErbB/EGFR signalling pathway was the most highly represented term, this identified EGFR signalling as putative regulatory mechanism of αVβ6 IACs. Indeed, based on these analyses, ErbB/EGFR signalling was more over-represented than adhesion signalling related terms (identified within the ARVC group). Moreover, many of the other top terms can also be associated with EGFR regulation or function (including cytokine-cytokine receptor interactions, MAPK signalling and endocytosis). Reactome pathway analysis also identified ErbB signalling as an overrepresented term, represented by the RAF/MAPK cascade group (Figure S4). Therefore, we confirmed the presence of EGFR in isolated αVβ6-dependent adhesion complexes (Figure S5A) and generated a protein-protein interaction network for all proteins in the complete dataset that were associated with the ErbB/EGFR signalling KEGG term and their one-hop protein interactors (to identify functionally linked proteins that did not meet the criteria of statistical significance or enrichment on LAP) (Figure S5B). Analysis of this network, and the αVβ6 IAC-specific dataset as a whole, enables identification and dissection of putative regulatory molecules of sub-networks that could regulate αVβ6-EGFR crosstalk.

### Phospho-proteomic analysis of αVβ6-dependent kinase signalling

An alternative approach that we employed to dissect αVβ6 signalling was to characterise signalling-associated phosphorylation events triggered by ligand-induced stimulation and endocytosis of αvβ6. BT-20 TNBC cells were stimulated with LAP as a soluble ligand (validation experiments showed that LAP stimulation triggered endocytosis of αvβ6 within 30 mins; data not shown). To assess temporal regulation of αvβ6-dependent kinase activity and phospho-signalling, a time-course of LAP stimulation was employed. Samples were phospho-enriched and subjected to analysis by mass spectrometry. To ensure the quality of the proteomic data, reproducibility across datasets was assessed principal component analysis employed to determine dataset-wide variance. For further details, see Supplementary Results Section: Quality assurance of phospho-proteomic analyses.

The phospho-proteomic dataset was interrogated using kinase substrate enrichment analysis (KSEA); a computational method ^24, 25^ to infer kinase activity based on identified phosphopeptides that are known substrates of specified kinase(s) moieties. For further details, see Supplementary Results Section: Kinase substrate enrichment analysis. The KSEA dataset was subject to hierarchical clustering to identify groups of kinases with similar activation/inactivation profiles (Clusters A-J). Individual kinase clusters, and groups of clusters, were then subjected to functional^26^ enrichment analysis. Over-representation of gene ontology (GO) term analysis was performed with KEGG (Kyoto Encyclopedia of Genes and Genomes) and Reactome pathway terms (Figure 4/S6C and S6C, respectively). ClueGO ontological term grouping was used to combine related terms into groups. Interestingly, KEGG analysis of “activated kinases” (clusters A-D) identified enrichment of proteins associated with ErbB/EGFR signalling as the most highly represented term (Figure 4A/B). Again, this identified EGFR signalling as a putative regulatory mechanism of αVβ6 function.

**Figure 4:**
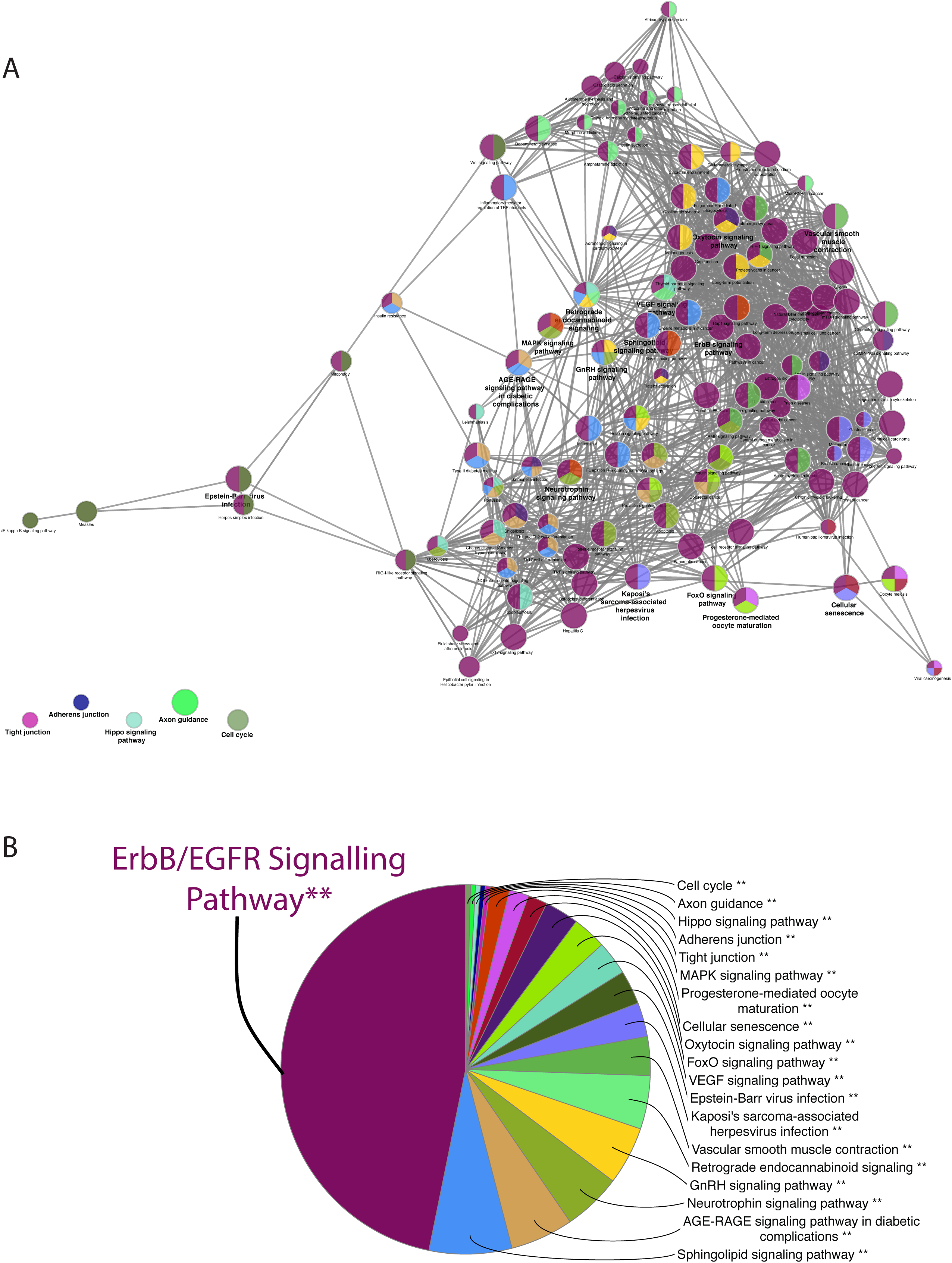
Kinase activation following ligand-dependent αVβ6 stimulation reveals αVβ6 EGFR crosstalk. Kinase Substrate Enrichment Analysis (KSEA) in BT-20 TNBC cells following stimulation of αVβ6 with soluble latency-associated peptide (LAP) (Time points: 0, 5, 15 & 30 mins), to infer kinase network plasticity during integrin αvβ6 LAP-engagement and internalisation ClueGO KEGG Pathway term hierarchical clustering. **A)** ClueGO hierarchical layout of KEGG terms identified for “activated kinases” (clusters A-D). Node colour corresponds to grouping. Nodes represent individual KEGG pathway terms. See supplementary Figure S6. **B)** Pie-chart organised by percentage of genes per term. Lead term highlighted by colour-coded text. Analysis parameters: minimum 2 genes per cluster, GO term/pathway network connectivity (Kappa score) =1; Statistical test Enrichment/Depletion (Two-sided hypergeometric test), Bonferroni p-value correction; GO term grouping based on kappa score, 50% of genes for group merge, 50% terms for group merge. Leading group term based on % gene/term.

### Integrin αVβ6-EGFR crosstalk regulates reciprocal receptor trafficking mechanisms

As KEGG analysis of αVβ6-dependent IACs (Figure 3 & S5]), and LAP-stimulated regulation of kinase activity (Figure 4 & S6), identified ErbB/EGFR signalling as a potentially key pathway regulated by αVβ6 (Figures 3 & 4), we assessed the subcellular distribution of both αVβ6 and EGFR. Consistent with identification of EGFR in αVβ6-dependent IACs (Figure 2C/D & S5A), under steady-state conditions in MDA-MB-468 cells, immunofluorescence demonstrated co-localisation between αVβ6 and EGFR in adhesion sites and also at intracellular structures reminiscent of trafficking vesicles (Figure 5A). To investigate this further, proximity ligation assays were performed and revealed that αVβ6 and EGFR co-exist in close proximity <40nm), suggesting they may form part of the same multi-molecular complex (Figure 5B). The distribution of the proximity ligation signal appeared to be predominantly intracellular. Since ontological analysis of αVβ6-dependent IACs, identified both ErbB/EGFR signalling and endocytosis as key pathways recruited to, or regulated by, αVβ6 (Figure 3), and αVβ6 and EGFR co-localised on intracellular structures (Figure 5A/B), we investigated whether EGFR and αVβ6 may regulate mutual trafficking pathways.

**Figure 5:**
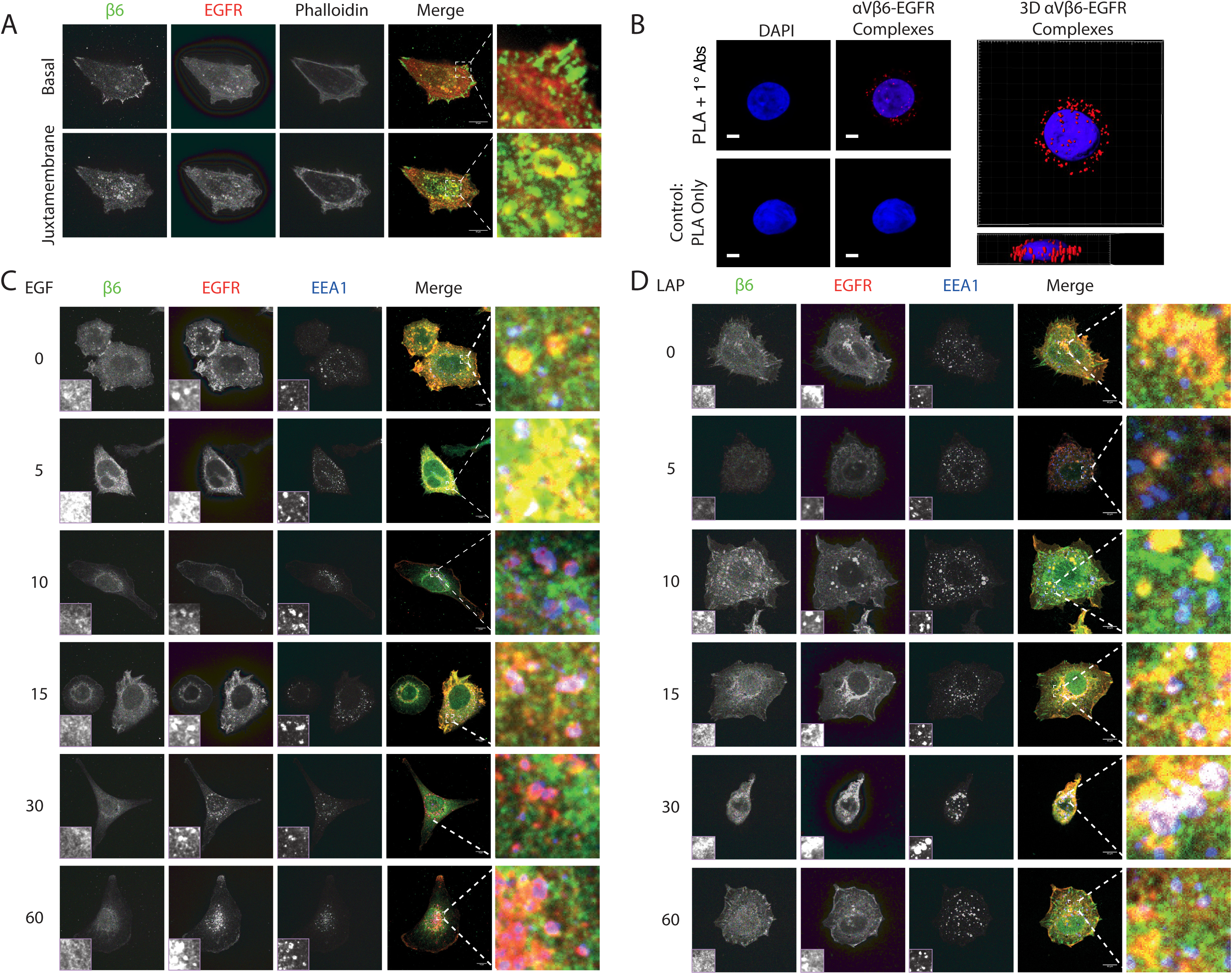
αVβ6-EGFR crosstalk regulates reciprocal receptor trafficking mechanisms. **A)**αVβ6 co-localises with EGFR in IACs and intracellular vesicles. Subcellular distribution of β6 and EGFR in MDA-MB-468 cells on glass coverslips in the presence of 10% FBS. Single z sections are displayed for basal and juxtramembrane (1 μm above basal) region. β6; green and EGFR; red. Scale bar = 10μm. **B)αVβ6 and EGFR are in close proximity.** Proximity ligation assay was performed using antibodies against αVβ6 (10D5, Millipore) and EGFR (D38B1, NEB) in MDA-MB-468. Positive signal confirmed that αVβ6 and EGFR epitopes are within 40nM of each other. Proximity signal is shown in red and the nucleus is stained blue (DAPI). Negative control contained PLA solution only. 3D representation of the localization observed in the first panels are shown in right-hand panel. Magnification bar = 5μM. **C)** EGF stimulation induces co-localisation of β6 and EGFR with EEA1. MDA-MB-468 cells, co-stained for β6, EGFR (D38B1) and EEA1. Single z slice shown for a juxtamembrane section of the cell. EGF stimulation is shown in minutes. for 10, 30 and 60 minutes, intensity arbitrary units. Scale bar = 10 μm. **D)** LAP stimulation induces alisation of β6 and EGFR with EEA1. MDA-MB-468 cells, co-stained for β6 (620W7), EGFR (D38B1) and EEA1. Single z slice shown for a juxtamembrane section of the cell. LAP stimulation is shown in minutes. Scale bar = 10 μm.

Under serum-starved conditions αVβ6 and EGFR co-localised, but not with EEA1 (early endosomal antigen 1; early endosome marker^26^), HRS (hepatocyte growth factor-regulated tyrosine kinase substrate; early endosome and multi-vesicular body marker^27, 28^) or LAMP2 (lysosome-associated membrane protein 2; lysosome marker^29^). However, EGF stimulation induced αVβ6 and EGFR co-accumulation on EEA1-positive vesicles between (10-30 mins post-stimulation), followed by a return to αVβ6 and EGFR co-localisation without EEA1 (60 mins post-stimulation) (Figure 5C). A similar pattern of co-localisation was observed with HRS, a marker of vesicles that are later in the endocytic trafficking pathway. Integrin αVβ6 and EGFR co-localised with HRS-positive vesicles after 15 mins EGF stimulation, however, unlike EEA1, a limited degree of αVβ6 and EGFR co-localisation with HRS persisted in all cells after 60 mins (Figure S7A). These data show that EGF induces re-distribution of αVβ6 in a manner consistent with internalisation and suggest that these receptors could have co-endocytosed together.

By contrast, in the presence of a combination of lysosomal and proteasomal inhibitors, EGF stimulation induced trafficking of EGFR to LAMP2-positive lysosomes (15-90 mins post-stimulation), whereas αVβ6 integrin was excluded from sites of EGFR and LAMP2 co-localisation (Figure S7B). Together these data suggest that EGF induces co-endocytosis of αVβ6 and EGFR, but that their trafficking pathways diverge after recruitment to HRS-positive endosomes; with EGFR being sent for lysosomal degradation and αVβ6 following an alternative trafficking route. These different fates for EGFR and αVβ6 agree with the canonical view that EGFR is targeted for lysosomal degradation, and integrins are predominantly recycled^30, 31^.

LAP stimulation was hypothesised to trigger endocytosis of αVβ6, as ligand stimulation is known to trigger integrin internalisation^30^. Therefore, a reciprocal experiment with LAP stimulation was performed to test whether soluble LAP stimulates αVβ6 endocytosis, and if so, whether αVβ6 internalisation affects the distribution of EGFR. LAP stimulation induced a degree of αVβ6 and EGFR co-localisation on EEA1-positive early endosomes in all cells (peaking at 10-15 mins post-stimulation) followed by a return to αVβ6 and EGFR co-localisation without EEA1 (30-60 mins post-stimulation) (Figure 5D). A similar pattern of co-localisation was observed with αVβ6, EGFR and HRS (Figure S7C). As observed for EGF stimulation, ligand-induced stimulation of αVβ6 with LAP induced trafficking of EGFR to LAMP2-positive lysosomes (30-90 mins post-stimulation), whereas αVβ6 did not co-localise with LAMP2 and is presumably targeted to alternative trafficking routes (Figure S7D).

Together these data indicate that ligand-induced endocytosis of either EGFR or αVβ6 triggers the internalisation of the other reciprocal receptor, which then traffic together to the MVB. Following this, EGFR is likely targeted for lysosomal degradation, whereas αVβ6 adopts an alternative pathway. This suggests mutual and reciprocally-regulated trafficking mechanisms for αVβ6 and EGFR.

### EGF stimulation suppresses αVβ6-mediated force-application on the matrix

Based on our proteomic, phospho-proteomic and imaging data, we reasoned that EGF-induced stimulation of αVβ6 endocytosis could serve as an EGFR-dependent mechanism to limit αVβ6 bioavailability and substrate engagement. Relative to other fibronectin-binding integrins, αVβ6 exhibits distinct biophysical properties that promote force-generation, rigidity sensing and increase matrix stiffening^15^. Moreover, TGFβ is mechanically activated by αVβ6; requiring transmission of intracellular actomyosin-dependent forces to be applied on latent TGFβ via αVβ6^32-34^. Biochemical integrin internalisation assays confirmed that EGF stimulation triggered endocytosis of αVβ6 (Figure 6A). Therefore, we investigated whether αVβ6-EGFR crosstalk could regulate the ability of αVβ6 to transmit cytoskeletal mechanical forces to the extracellular environment. We performed traction force microscopy, using polyacrylamide hydrogels coated with the αVβ6-selective ligand LAP, to measure αVβ6-dependent mechanical force application^32^. These assays revealed that stimulation with EGF induced a marked suppression in the ability of MDA-MB-468 cells to apply αVβ6-dependent mechanical force on LAP (Figure 6B). These data suggest that, consistent with EGF inducing endocytosis of αVβ6 (Figure 6A), EGFR stimulation limits the availability of αVβ6 at the cell surface to apply extracellular mechanical forces.

**Figure 6:**
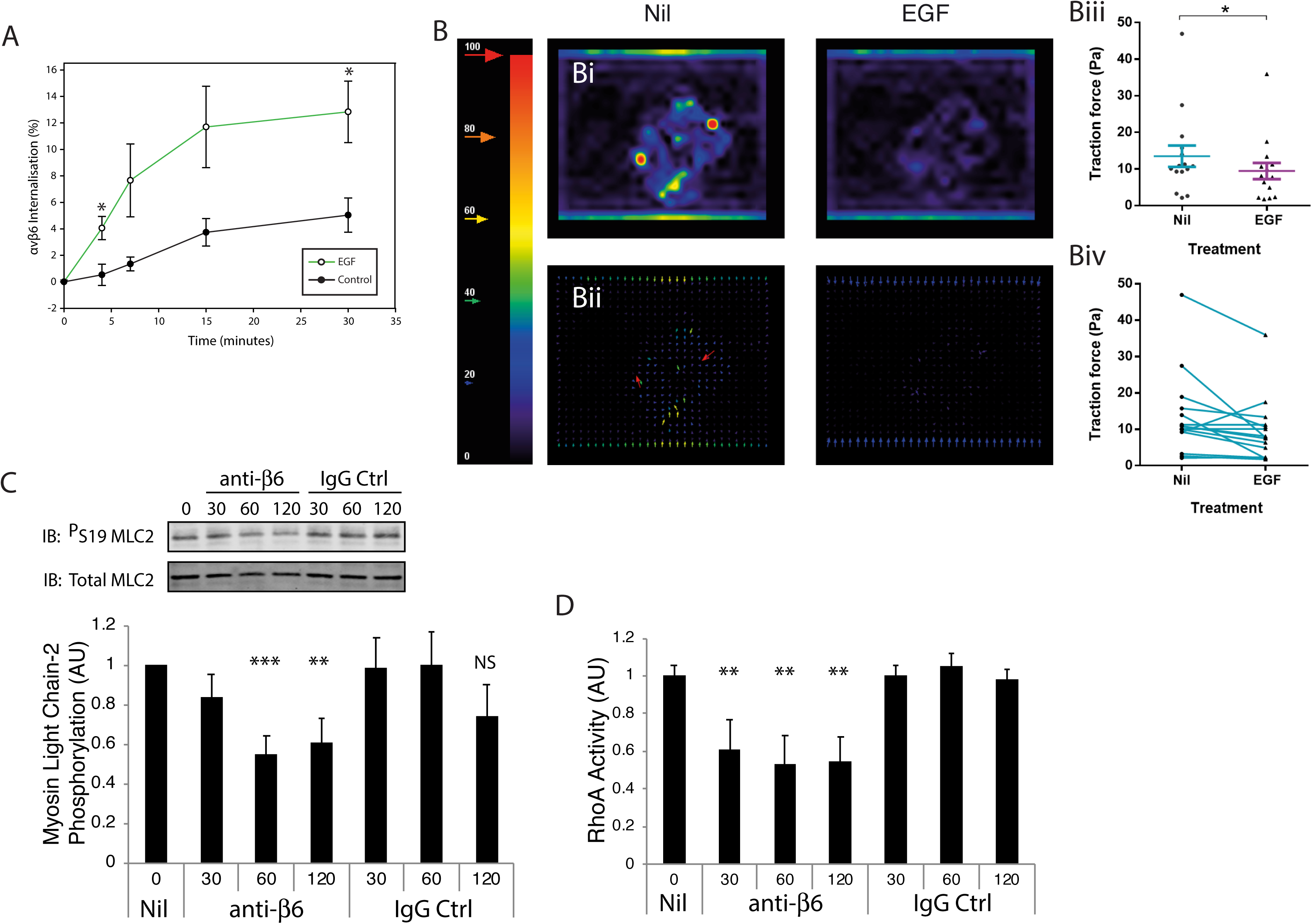
EGF stimulation induces αVβ6 endocytosis and suppresses αVβ6-mediated force-application. **A)** EGF stimulates αVβ6 endocytosis. Integrin endocytosis assay showing percentage αVβ6 internalisation at 4, 7, 15 and 30 minutes with EGF stimulation (10 ng/ml) or vehicle control (0.5 μg/ml). Receptor internalisation is expressed as a percentage of the total surface levels on unstimulated cells. Mean and SEM of data points is plotted. One-way ANOVA * = p< 0.05, N = 3. **B)** EGF stimulation supresses αVβ6-mediated force-application on LAP. Traction force microscopy was performed on MDA-MB-468 cells were cultured on LAP-coated 0.4kPa polyacrylamide hydrogels. MIcrospheres and cells were imaged before and after addition of EGF. After analysis, traction force values for each cell were calculated for before (Nil) and after EGF treatment (EGF). Force magnitude **(Bi)** and vector **(Bii)** plots are shown for representative cells. Graphs show either the cumulative data of the cell population **(Biii)** or the effect of EGF stimulation on traction forces applied by individual cells (blue lines show change in force application following EGF stimulation for each cell) **(Biv)**. Error bars = S.D. *P < 0.05. **C)** Antibody blockade of αVβ6 suppresses MLC2 phosphorylation. MDA-MB-468 cells, pre-spread on FN, were treated with anti-αVβ6 blocking antibody 53A2 (10μg/ml) over a time-course (0, 30, 60 and 120 mins). Phosphorylation of MLC2 (^P^S19) was assessed by immunoblotting and normal to total MLC2 protein levels. N=4, error bars = SEM. **D)** αVβ6 inhibition reduces RhoA activity. MDA-MB-468 cells, pre-spread on FN, were treated with anti-αVβ6 blocking antibody 53A2 (10μg/ml) over a time-course (0, 30, 60 and 120 mins). RhoA activity was assessed by G-LISA. N=4, error bars = SEM.

Given the potential role of αVβ6 in force transduction mechanisms, the capacity of αVβ6-EGFR crosstalk to modulate the ability of cells to apply αVβ6-dependent forces on the extracellular microenvironment could have a direct impact on matrix rigidity sensing, ECM remodelling and TGFβ activation^15, 34^; suggesting a mechanism by which mechanotransduction and mechanosensation could be fine-tuned spatially and temporally.

### αVβ6 Integrin regulates nuclear shuttling of force-dependent transcriptional co-activators

Having identified a mechanism by which αVβ6 regulates extracellular transmission of mechanical forces, we assessed the impact of αVβ6 on transmission of intracellular force. In addition to identification of the ErbB/EGFR signalling pathway (Figure 3 & S5), KEGG functional enrichment analysis identified over-representation of Hippo pathway regulators at αVβ6-dependent IACs (Figure 3 & S8). The Hippo signalling pathway represents a phosphorylation cascade that regulates the transcriptional activators YAP/TAZ. Phosphorylation of YAP/TAZ prevents their translocation to the nucleus and subsequent stimulation of proliferative and anti-apoptotic gene transcription^35^. Mechanosignalling is a key regulator of YAP/TAZ nuclear translocation; mechanical inputs such as stiff ECM and cytoskeletal tension drive YAP/TAZ to accumulate in the nucleus to activate transcription of target genes^36-38^.

Therefore, we sought to determine whether αVβ6 regulates the force-dependent transcription co-activators YAP. First, we investigated the ability of αVβ6 to modulate cytoskeletal contractility. Importantly, immunoblotting revealed that antibody-mediated blockade of αVβ6 inhibited Myosin Light Chain-2 phosphorylation (MLC2 ^P^S19); a key marker of actomyosin contractility (Fig. 6C). Moreover, G-LISA-based biochemical analysis of the small GTPase RhoA, a key regulator of actomyosin contractility, demonstrated that αVβ6 inhibition suppressed RhoA activity levels (Fig. 6D). Together, these data suggest that αVβ6 plays a major role in cytoskeletal regulation and actomyosin-dependent force transmission in MDA-MB-468 cells. Therefore, we examined the role of αVβ6 in regulating nuclear shuttling of YAP on 2D stiff substrates (FN-coated glass) and in 3D collagen gels embedded with FN and LAP of two different rigidities (Figure 7). As expected, given the force-sensitive nature of YAP translocation, the number of cells with entirely nuclear YAP distribution is reduced in compliant 3D matrices, compared to a stiff 2D substrates. Under control conditions, the majority of cells on stiff 2D substrates demonstrated exclusively nuclear YAP distribution (Figure 7Ci), whereas control cells in high rigidity 3D matrices (3mg/ml collagen + FN/LAP) exhibited nucleo-cytoplasmic YAP (Figure 7A/B/Cii). However, consistent with the ability of αVβ6 to regulate mechanotransduction and actomyosin contractility, antibody-mediated inhibition of αVβ6 substantially reduced nuclear localisation of YAP. On stiff 2D substrates αVβ6 blockade induced nucleo-cytoplasmic YAP distribution (Figure Ci), whereas in high rigidity 3D matrices (3mg/ml collagen + FN/LAP) αVβ6 inhibition excluded YAP from the nucleus (Figure 7A/Cii); highlighting the critical role that αVβ6 plays in regulating force-sensitive transcription factor function. Similar, but supressed, responses were observed in softer 3D matrices (1.7mg/ml collagen + FN/LAP), in which. cells exhibited lower levels of baseline nucleo-cytoplasmic distribution under control conditions (Fig 2Ciii). Together, these data suggest that αVβ6 blockade suppresses force-transmission and dysregulates YAP localisation and transcriptional reprogramming.

**Figure 7:**
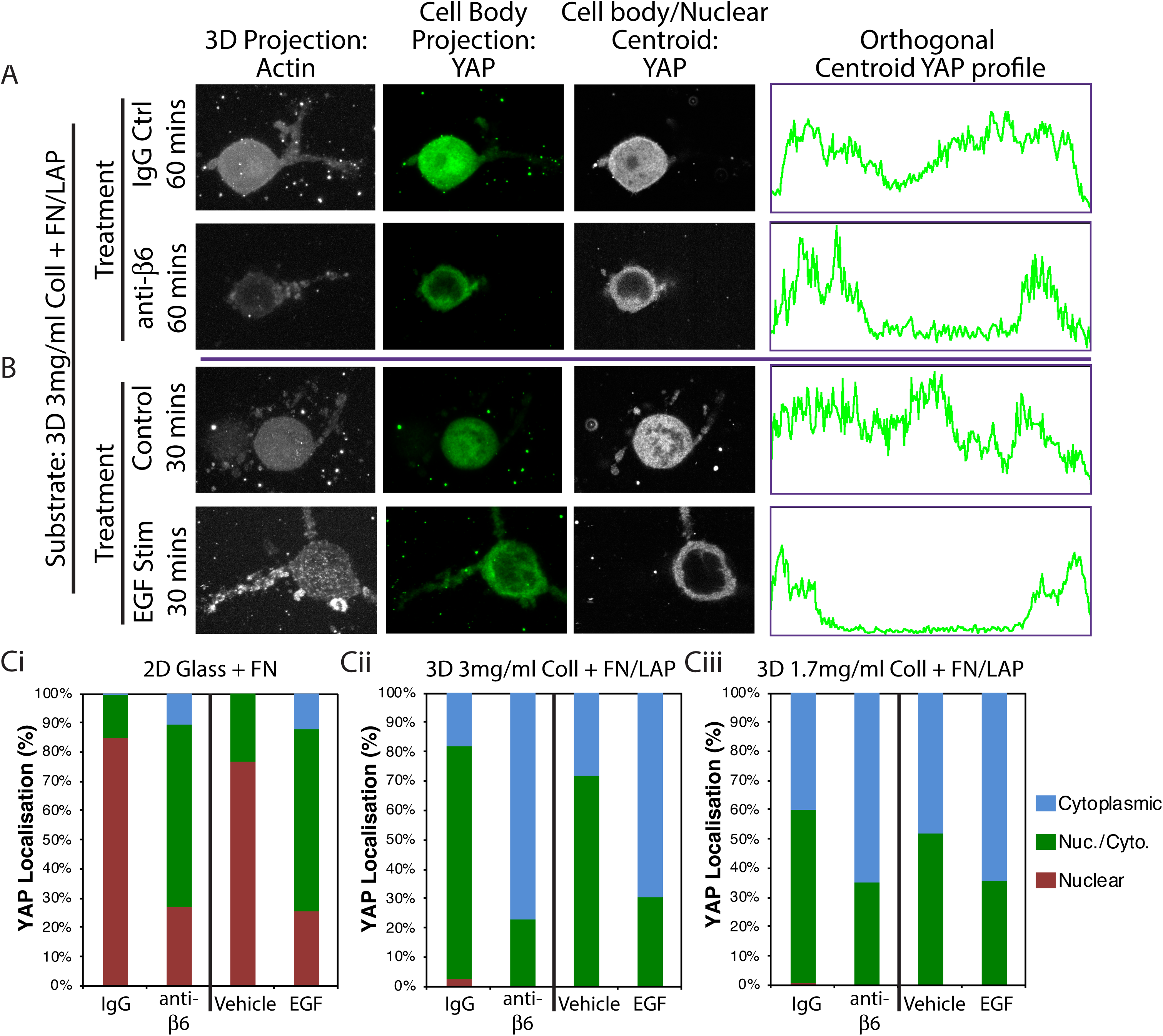
αVβ6 Integrin and EGF regulate nuclear shuttling of force-dependent transcriptional co-activators. **A & B)** Immunofluorescence micrographs showing subcellular localisation of YAP in MDA-MB-468 cells in “high rigidity 3D matrices” (3mg/ml 3D collagen gels supplemented with 25 μg/ml FN and 0.5 μg/ml LAP). ”3D Projection: Actin”: Maximum projection of z-sections of phalloidin staining for entire 3D volume of cells. ”Cell Body Projection: YAP”: Sum projections of z-sections of YAP staining through the volume of the cell body; All projections represent the same 3D volumes. ”Cell body/Nuclear Centroid: YAP”: Single z-section of YAP staining is displayed representing the centre of the nucleus. ”Orthogonal Centroid YAP Profile”: Line profile displaying YAP signal intensity for a single line ROI across the centroid orthogonal plane of the nucleus> A) Cells seeded in 3D gels in presence of 10% FBS, and serum-starved for 2.5 hrs prior to 60 mins antibody treatment (10 μg/ml Rat IgG or anti-αVβ6 blocking antibody (620W7)). Cells seeded in 3D gels in presence of 10% FBS, and serum-starved for 3 hrs prior to 30 mins treatment (Vehicle control or 10 ng/ml EGF). Visual scoring of YAP distribution: nuclear (Nuclear), nucleo-cytoplasmic (Nuc./Cyto.) and cytoplasmic (Cytoplasmic) distribution. Cells seeded on different substrates: Ci) “Stiff 2D substrate” (glass coverslip coated with 10 μg/ml FN); Cii) “high rigidity 3D matrices” (3mg/ml 3D collagen gels supplemented with 25 μg/ml FN and 0.5 μg/ml LAP); “softer 3D matrices” (1.7mg/ml 3D collagen gels supplemented with 25 μg/ml FN and 0.5 μg/ml LAP. Data representative of 3 independent experiments

Having established a role for αVβ6-EGFR crosstalk in regulation of αVβ6-dependent mechanical forces (Figure 6A/B), we assessed the impact of EGF stimulation on YAP localisation in MDA-MB-468 cells. Interestingly, on 2D substrates and in 3D matrices, EGF stimulation recapitulated the effect of αVβ6 inhibition (Figure 7B/Ci/Cii/Ciii). This observation is consistent with the role of EGF in stimulating αVβ6 endocytosis, to limit αVβ6-ECM engagement and mechanotransduction (Figures 5 & 6). However, to date, we cannot discount that this effect of EGF on YAP distribution is independent of αVβ6 function.

### αVβ6 and EGFR co-operate to drive TNBC invasion

Having identified a functional relationship between αVβ6 and EGFR, that impacts receptor trafficking, ligand-engagement and bi-directional force transduction, we assessed whether αVβ6 and EGFR gene expression correlated in patient tissue. Analysis of the METABRIC cohort showed that *EGFR* mRNA profiles exhibited a trend of significant positive correlation with *ITGB6* (rho=0·14, *P*=1·63 × 10^-10^). (Figure S1G). Thus, we considered that αVβ6-EGFR crosstalk might influence the pro-invasive activities in TNBC cells.

We analysed invasion in a panel of three αVβ6- and EGFR-positive TNBC cell lines (HCC38, MDA-MB-468 and BT-20). siRNA-mediated inhibition of ITGB6 (β6 subunit) or EGFR, or blockade of EGFR activity (using gefitinib, GEF) significantly blocked invasion in all cell lines (Figures 2A-C), thus confirming invasion was αVβ6- and EGFR-dependent.

Based on our previous data on αVβ6-EGFR crosstalk, we further considered that co-regulation of αVβ6 and EGFR activity might influence pro-invasive activities. Therefore, EGF was added to cells to induce EGFR activation. EGF significantly increased the invasive capacity of MDA-MB-468 and BT-20 cells, but the high baseline level of HCC38 invasion was not further potentiated with EGF (Figure 8D-F). In all TNBC cells tested, the EGF-induced invasion was inhibited by siRNA-mediated knockdown of ITGB6 or EGFR (Figure 3), or by blockade of EGFR (Gefitinib, GEF).

**Figure 8:**
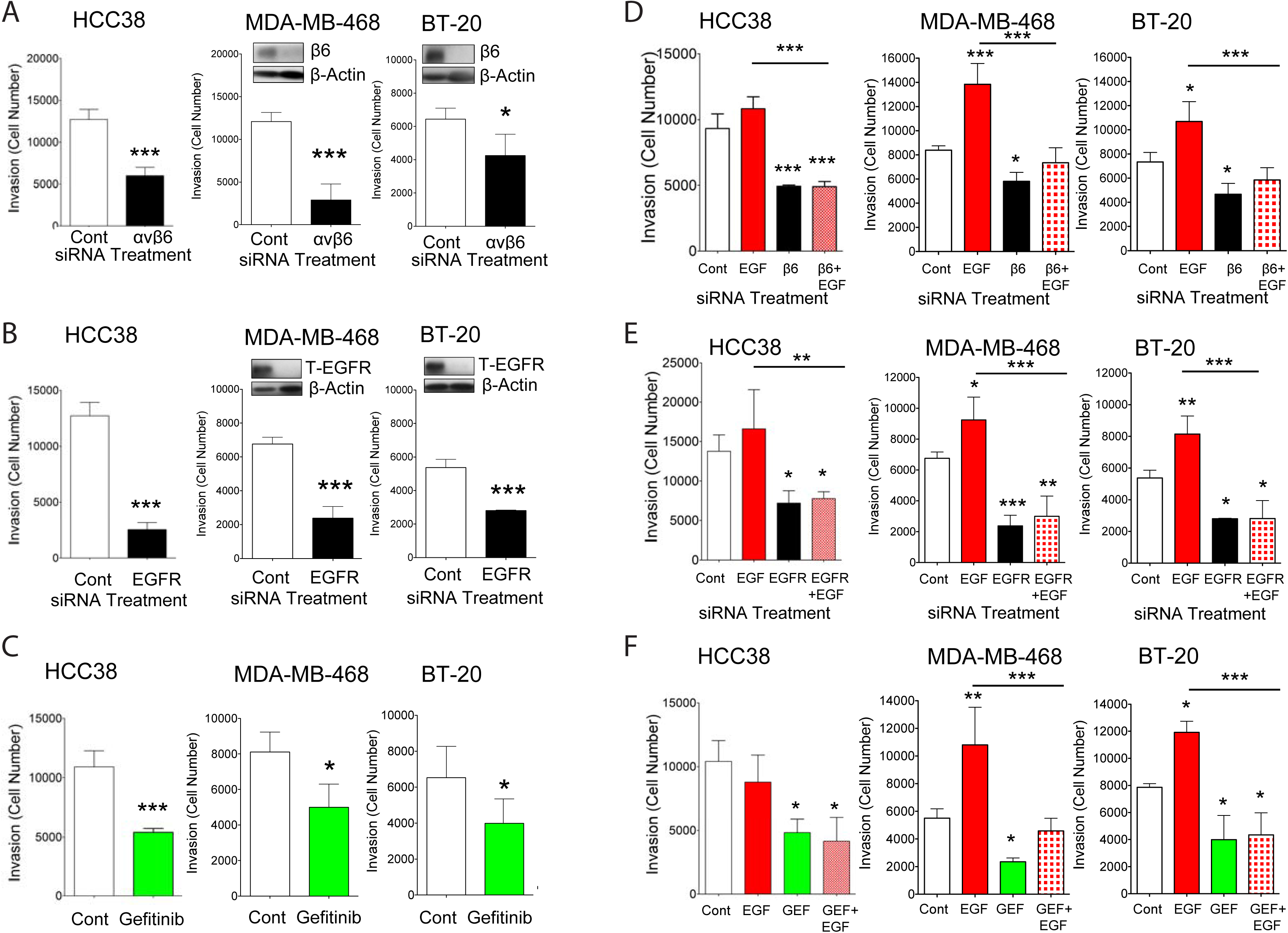
αVβ6 regulates EGFR-driven TNBC cell invasion. Transwell invasion assay of HCC38, MDA-MB-468 and BT-20 TNBC breast cancer cell lines which all express integrin αVβ6 and EGFR. 5 × 10^4^ cells/well were seeded and the number of cells that invaded was counted after 72h. **A)** Breast cancer cell line invasion is integrin αVβ6-dependent. Transwell invasion of cells transfected with control or β6 siRNA (20nM). **B & C)** Breast cancer cell line invasion is EGFR-dependent. Transwell invasion of cells treated with IgG or EGFR inhibitor Gefitinib (GEF) (10μg/ml) **(B)** or transfected with control or EGFR siRNA (20nM) **(C)**. **D & E)** EGF stimulated breast cancer cell line invasion is αVβ6- and EGFR-dependent. HCC38, MDA- MB-468 and BT-20 TNBC cells transfected with control, β6 **(D)** or EGFR **(E)** siRNA (20nM), were pre-treated for 30 min with vehicle or EGF (1 × 10^-5^M) prior to seeding in Transwell invasion assays. **F)** Transwell invasion of cells pre-treated with IgG or Gefitinib (GEF) (10μg/ml) in the presence or absence of EGF (1x 10^-5^M). All experiments were performed in triplicate, representative experiments shown (N=4-6, error bars represent 95% Confidence Interval). \**P*=0·05, \*\**P*=0·01, \*\*\**P*<0·001 (relative to IgG or control treated cells).

These data suggest that EGFR-driven invasion is mediated by αVβ6. Moreover, the data predict that as αVβ6 regulates intracellular signals that may be required for EGFR-dependent invasion then αVβ6- blockade could enhance EGFR-targeted therapy.

## DISCUSSION

In this study, we investigated the role of αVβ6 integrin in TNBC. We report an association between high expression of αVβ6 and poor survival in both triple-negative breast cancer (TNBC) and ER-negative breast cancer, the worst prognostic sub-groups of breast cancer. To understand how αVβ6-dependent signalling mechanisms might contribute to TNBC, we employed novel proteomic and phospho-proteomic approaches to dissect αVβ6 signalling. These approaches provided a global view of αVβ6-mediated signalling and identified epidermal growth factor receptor (EGFR) signalling as a key regulatory pathway associated with αVβ6 function.

Further analysis revealed:

- Crosstalk between αVβ6 and EGFR regulates trafficking and engagement of each receptor
- αVβ6-EGFR crosstalk regulates αVβ6-mediated force-application on the matrix
- αVβ6 controls force-dependent nuclear localisation of transcriptional co-activator YAP
- αVβ6 and EGFR expression co-associate in breast cancer cells
- αVβ6 and EGFR co-operate to drive TNBC invasion

To understand how αVβ6 contributes to TNBC development, we employed two unbiased approaches to dissect αVβ6-dependent signalling mechanisms. First, we performed proteomic analysis on ligand-engaged αVβ6 IACs, to define the signalling networks recruited to sites of αVβ6-ECM interaction. Second, we used a phospho-proteomic strategy to identify kinase activation pathways that were activated following ligand-induced endocytosis of αVβ6. Both of these approaches, and follow-up analyses, identified the pro-invasive receptor tyrosine kinase EGFR as a key regulator of αVβ6 function.

Relative to other integrins, αVβ6 exhibits distinct biophysical properties that promote force-generation and increase matrix rigidity^15^, suggesting that αVβ6 could play a key role in matrix stiffening in breast cancer. However, our data suggest that a complex regulatory mechanism exists involving crosstalk between αVβ6 and the EGFR, which impacts mutual receptor trafficking mechanisms. Stimulation of EGFR triggers internalisation of αVβ6, and *vice versa*, thus limiting the bioavailability of αVβ6 integrin and inhibiting the ability of breast cancer cells to exert force on the ECM (Figure 6). Moreover, proteomic and imaging data demonstrated that EGFR is specifically recruited to sites of αVβ6-matrix engagement (Figures 2, 3 & S5), suggesting that engagement of αVβ6 at the cell-matrix interface recruits EGFR, whereas EGF triggers endocytosis of both αVβ6 and EGFR to reduce levels at the cell surface and suppress matrix stiffening (Figure 6A/B). Consistent with this mutually regulatory role, EGF-dependent invasion of a panel of triple-negative breast cancer cells requires αVβ6 activity (Figure 8).

These data suggest a model whereby αVβ6-EGFR crosstalk regulates matrix stiffening, but also the transmission of extracellular matrix forces into the cell, in order to co-ordinate transcriptional reprogramming and invasion. Such a mechanism has potential for positive feedback, whereby αVβ6-dependent matrix stiffening drives αVβ6-mediated intracellular force transduction. However, by controlling receptor trafficking, EGF-dependent regulation of αVβ6 would provide a mechanism by which bidirectional force transduction could be fine-tuned to spatially and temporally co-ordinate mechanotransduction and mechanosensation.

While αVβ6 has the capacity to promote force-generation and sense matrix rigidity^15^, αVβ6 also employs forces to activate TGFβ. Integrins αVβ6 and αVβ8 have the unique ability to activate TGFβ by applying force to induce a conformational change in LAP, thus releasing active TGFβ from its inhibitory complex^34, 39^. Indeed, αVβ6 has evolved structural specialisations that dictate a ligand-binding orientation that specifically supports force application and cytoskeletal tensile force transmission^33^. Therefore, while we haven’t addressed it explicitly in this study, it is likely that this regulatory mechanism could also impact αVβ6-dependent TGFβ activation. Indeed, in pancreatic cancer cells, EGF has been shown to modulate GTPase activity to control αVβ6-dependent migration, in a manner that opposes TGFβ activation^32^.

Integrin-mediated intracellular transmission of forces from ECM can modulate cell behaviour directly, through changes in adhesion signalling networks established at sites of integrin engagement (the adhesome), and indirectly, via force-dependent regulation of transcriptional processes. By controlling availability of αVβ6 at the cell-matrix interface, our data suggest that αVβ6-EGFR crosstalk could impact both of these mechanisms. YAP/TAZ are transcriptional regulators that shuttle between the cytoplasm and nucleus, where they modulate transcription, in a matrix stiffness- and force-dependent manner, to drive proliferation, invasion, epithelial to mesenchymal transition and metastasis^40^ YAP/TAZ accumulate in the nucleus in response to mechanical force and biochemical inputs and relay these signals by activating transcription of target genes ^36^. We have shown that inhibition of αVβ6 reduced YAP nuclear shuttling on 2D, and in 3D, matrices. Inhibition of αVβ6 also decreased RhoA activity and myosin-light chain phosphorylation. Together, these data suggest that αVβ6 blockade suppresses force-transmission and dysregulates YAP localisation and transcriptional reprogramming.

It is becoming increasingly clear that YAP/TAZ plays a key role in breast cancer progression. Translocation of YAP/TAZ to the nucleus is regulated by ECM rigidity, which is key prognostic indicator in breast cancer and driver of breast cancer invasion^1, 41-43^ and transcriptional reprogramming as a result of nuclear YAP/TAZ localisation leads poorer prognoses ^36, 44^. YAP/TAZ activity correlates with histological grade and metastases of breast cancer ^45^. TAZ nuclear expression is strongly associated with TNBC in comparison to ER/PG receptor-positive and ErbB2-positive breast cancer, and correlates with poorer clinical outcomes of recurrence and overall survival ^44^. Therefore, the ability of αVβ6 integrin to regulate YAP nuclear translocation may have direct clinical relevance in TNBC. Importantly, EGF stimulation also reduced nuclear YAP localisation (Figure 7). Given the relationship that we have described between αVβ6 and EGFR, it is possible that this is a consequence of triggering αVβ6 endocytosis and limiting αVβ6 levels at the cell-matrix interface. However, to date, we cannot discount the possibility that EGF may be exerting a direct effect on YAP distribution that is independent of αVβ6 function.

In this study we employed novel proteomic and phospho-proteomic techniques to dissect adhesion receptor signalling. These approaches enabled a global and dynamic view of αVβ6-mediated signalling to be established. This work represents the first attempt to define αVβ6-mediated signalling networks and, while they retained the same structural core, αVβ6-IACs were strikingly different from previously published IACs (which were largely focused on α5β1 and αVβ3 integrins binding FN)^6, 46-52^. Compared to other integrins, αVβ6 is a relatively under-studied integrin. Therefore, we anticipate that these datasets will provide a unique resource for studying the function of this clinically-important integrin. However, in future it will be necessary to compare αVβ6 signalling networks in different tumour types, which perhaps rely on different RTKs, in order to define the αVβ6-specific consensus and meta-adhesomes.

This is the first study to show an association between high αVβ6 expression in ER-negative and TNBC and poor survival, and further, that this occurs predominantly in younger women, a group that desperately needs novel effective therapies. Our data support the proposal that testing of biopsies for αVβ6 expression should become a routine immunopathological procedure to stratify patients with breast cancer into this new ‘very high’ risk αVβ6-positive/TNBC subgroup. In the longer term, this stratification may provide a therapeutic opportunity for this high-risk subgroup.

It will now be essential to assess the consequence of co-targeting αVβ6 and EGFR in TNBC tumour models. Our previous study showed that targeting αVβ6 with 264RAD reduced growth of HER2+ breast cancer xenografts *in vivo* (9), whereas, co-targeting both αVβ6 and HER2, with trastuzumab, was more effective than monotherapy against either target^8^. Targeting EGFR in breast cancer as a monotherapy with gefitinib has proved disappointing, with improved response limited to subsets of ER-positive breast cancer such as those with tamoxifen-resistance^53^; hence trials have progressed in combination with anti-hormonal therapies^54^. Trials of the therapeutic efficacy of gefitinib in TNBC (NCT01732276) are in progress. However, our *in vitro* data suggest that αVβ6 and EGFR exist in a multi-molecular complex and exhibit co-regulatory mechanisms that impact receptor availability, function and TNBC invasion. Thus it is clear that αVβ6 has the potential to regulate the therapeutic response to gefitinib, and *vice versa*.

However, to fully exploit adhesion receptors and receptor tyrosine kinases therapeutically, it is essential to understand the integration of their signalling functions and how crosstalk mechanisms influence invasion and the response of tumours to molecular therapeutics. Ultimately, the success of an αVβ6 therapeutic is likely to be highly-dependent on precise patient stratification. To fully predict patient responses to αVβ6-targeting drugs, and combination therapies, it will be necessary to employ systems-level analyses to understand the impact of key regulatory mechanisms on the proteomic, phospho-proteomic and transcriptional landscapes of both patient-derived tumour and stromal cells.

## MATERIALS AND METHODS

### Clinical Samples and Immunohistochemical Analysis

Two independent cohorts of breast cancer samples were analyzed following REMARK guidelines^55^. Immunohistochemistry utilized 4μm, formalin-fixed, paraffin-embedded serial sections of tissue microarrays (TMA). The protocol used for αVβ6 integrin (mAb 6.2G2,Biogen Idec) was described previously^9^.

### METABRIC cohort pre-processing

This study makes use of the METABRIC data generated by the Molecular Taxonomy of Breast Cancer International Consortium^56^. Funding for the project was provided by Cancer Research UK and the British Columbia Cancer Agency Branch. Breast cancer METABRIC dataset was preprocessed, summarized and quantile-normalized from the raw expression files generated by Illumina BeadStudio. (R packages: beadarray v2.4.2 and illuminaHuman v3.db_1.12.2). Raw METABRIC files were downloaded from European genome-phenome archive (EGA) (study id: EGAS00000000083). Raw data files of one METABRIC sample was not available at the time of our analysis, therefore it was excluded. All preprocessing was performed in R statistical environment v2.14.1.

### Statistical Analysis of Clinical Data

London and Nottingham clinical cohort risk groups (low expression and high expression) were established by dichotomizing αVβ6 protein abundance using the thresholds derived from the previously published IHC dataset on αVβ6^8^. Consistent with Moore et al (2014), the London subset ER and PR thresholds were <3 (negative) and ≥3 (positive). However, very low HER2 expression was treated as negative in this study, i.e. where HER2<3. In the Nottingham subset, positivity and negativity were pre-assigned and hence used as is.

London and Nottingham datasets were dichotomised into low- and high-risk groups using αVβ6 protein expression (Low-risk αVβ6<5, High-risk αVβ6≥5). Survival analysis was performed in R statistical environment v.2.14.1 (R package:survival v2.36-14). Hazard ratio was estimated by fitting univariate Cox proportional hazards model, and significance of difference between the survival of risk groups were computed using Logrank test. For the METABRIC cohort^56^, risk groups (low expression and high expression) were established by dichotimising ITGB6 mRNA abundance using median expression. For METABRIC published data, ER (IHC calls +/-), PR (gene expression based +/-calls), and HER2 (IHC calls 0,1,2,3) were used to determine triple receptor negative patients. HER2 staining scores of 0, 1 and 2 were classed as negative.

### Proteomic analysis of αVβ6 integrin-associated complexes

#### - Isolation of Integrin-Associated Complexes

The protocol for isolating integrin-associated adhesion complexes is based on a published methodology with some outlined modifications ^57^. Tissue culture 10 cm dishes were coated with either fibronectin, LAP or collagen I ligands overnight at 4°C, washed twice in PBS (-), and blocked using heat-inactivated BSA for 30 minutes at room temperature. Plates were washed twice in PBS (-), once in DMEM/25mM HEPES, then incubated with 9 ml of DMEM/25mM HEPES to equilibrate at 37°C, 5% CO_2_.

Cells were washed in PBS (-), harvested by trypsinisation and centrifuged at 280 × g for 4 minutes. Cell pellets were then re-suspended in DMEM/25mM HEPES and incubated in 40 ml DMEM/25mM HEPES at 37°C for 30 minutes. The cell suspension was centrifuged and re-suspended in DMEM/25mM HEPES sufficient to plate 1 ml of cell suspension per prepared 10 cm dish. Cells were seeded at 5 × 106 per ml and allowed to adhere to ligand for 2 hours 30 minutes at 37°C, 5% CO_2_.

Cells were then cross-linked with the cell-permeable crosslinker DTBP (dimethyl 3,3’-dithiobispropionimidate), which stabilises protein interactions. DMEM/25mM HEPES was then removed and replaced with 5 ml per plate of pre-warmed (37°C) 3 mM DTBP in DMEM/25mM HEPES. Plates are incubated at 37°C for 30 minutes to permit cross-linking.

Cells were then washed twice in ice cold PBS (-) and incubated with 20 mM pH 8.0 Tris for 5 minutes at room temperature, to quench the crosslinker activity. Plates were then washed twice in ice cold PBS (-) and transferred to ice packs.

Immediately prior to sonication, cells were washed once and filled with cold extraction buffer (20 mM NH_4_OH, 0.5% (v/v) Triton X-100 in PBS (-)). Cells were sonicated submerged in extraction buffer, using the SONICS Vibra cell ™ sonicator at 20% amplitude for approximately 2 minutes per plate. After sonication plates were washed three times in cold extraction buffer, and three times in cold PBS(-).

PBS (-) was then thoroughly removed from the plates, and plates were dried. Remaining adhesion complexes were harvested by scraping in 2 × concentrated reducing Sodium dodecyl sulfate (SDS) sample buffer. Harvested material was collected in an Eppendorf and incubated at 95°C for 5 minutes. A proportion of the sample was then immunoblotted to assess the quality and specificity of the adhesion isolation. The remaining sample was then processed for MS.

### Sample Preparation for Mass Spectrometry

Samples from 2D cell-matrix adhesion isolation preparation were resolved by polyacrylamide gel electrophoresis (PAGE), in 1.5 mm 10 well 4 – 12% Novex® NuPAGE™ gels (Invitrogen), in the Novex® Mini-cell XCell SureLock™ Electrophoresis tanks (Invitrogen). A constant 160 V was used to resolve the samples. Novex® NuPAGE™ MES-SDS running buffer (Invitrogen) was used.

Protein in the gels was stained by incubating the gel in Instant Blue™ (Expedion) colloidal Coomassie protein stain, for one hour at room temperature on the rocker. Gels were then de-stained with five five-minute washes in MilliQ water, on the rocker. Gels were then washed further in MilliQ for one hour on the rocker, before storing in MilliQ overnight at 4°C.

Gel lanes were excised with sterile scalpel blades on a clean tile. Gel bands were then cut into ∼1 mm^3^ pieces and transferred into a single corresponding well of a 96-well perforated plate (Glygen Corp). The gel was kept moist throughout excision with MilliQ water. Gel pieces were then destained by incubation in 100 µl 50% (v/v) acetonitrile (ACN) / 50% (v/v) NH_4_HCO_3_ for 30 minutes at room temperature, before removing the supernatant (wash) by centrifugation (96-well-plate rotor, 1,500 RPM, 2 minutes). This step was repeated until all the stain was removed from the gel pieces, leaving them transparent (usually four washes were required).

Gel pieces were dehydrated twice by incubating with 50 µl 100% ACN for 5 minutes, removing the supernatant following centrifugation each time. Gel pieces were then dried by vacuum centrifugation for 20 minutes using the Christ VWR RVC 2-25 Speed vacuum, then incubated in 50 µl 10 Mm Dithiothreitol (DTT) for one hour at 56°C, to reduce the proteins in the sample. DTT was then removed following centrifugation, and proteins then alkylated by incubating the gel pieces in 50 µl 55 mM Iodoacetamide (IA) for 45 minutes at room temperature, in the dark.

Following IA removal, gel pieces were sequentially washed and dehydrated twice, by incubating in 50 µl 25 mM NH_4_HCO_3_ for 10 minutes, then 50 µl ACN for 5 minutes at room temperature, removing each solution after incubation following centrifugation. The gel pieces were then dried by vacuum centrifugation for 20 minutes prior to incubation with 1.25 ng/µl trypsin gold (Promega) in 25 mM NH_4_HCO_3_ for 45 minutes at 4°C and overnight at 37°C.

Tryptic peptides were collected by centrifugation. Additional peptides were extracted by incubating the gel pieces in 50 μl 99.8% (v/v) ACN/ 0.2% (v/v) formic acid (FA) for 30 minutes at room temperature, centrifugation, followed by incubating with 50 μl 50% (v/v) ACN/ 0.1% (v/v) FA for 30 minutes at room temperature. These additional extracted peptides were collected by centrifugation then pooled with the initial supernatant and evaporated to dryness in the collection plate by vacuum centrifugation. Dried peptides were re-suspended in 20 µl 5% (v/v) ACN in 0.1% FA and stored at –20°C until analysis.

### Mass Spectrometry

4µl of each digested fraction was injected onto a Nanoacquity™ (Waters) Ultra Performance Liquid Chromatography (UPLC) column, coupled to an LTQ-Orbitrap XL (Thermo Fisher) equipped with a nanoelectrospray source (Proxeon). Samples were separated on a 1 – 85% ACN gradient, 0.300 µl/min flow rate, with an 80-minute retention time. Dynamic exclusion was enabled for a repeat count of 1 for a duration of 30.00 s. MS spectra were acquired by the LTQ-Orbitrap at a resolution of 30,000 and MS/MS was performed on the top 12 most intense ions in the LTQ ion trap.

### Peptide Identification and Proteomic Analysis

Raw peptide MS data were converted into peak lists and searched against a reviewed H. Sapiens UniProt database (containing 149,633 sequences; 47,132,354 residues) using Mascot Daemon (version 2.3.2) software. The initial precursor and fragment ion maximum mass deviations in the database search were set to 5 ppm and 0.6 Da, respectively, which is optimal for linear ion trap data. One missed cleavage by the enzyme trypsin was allowed. Cysteine carbamidomethylation (C) was set as a fixed modification, whereas oxidation (M, K, P), and phosphorylation (S, T, Y) were considered as variable modifications.

The database search results were processed and statistically evaluated within Scaffold (version 4) The false discovery rate (FDR) for the peptides and proteins were set to 0.01 and 0.4 respectively, to ensure the worst peptide/protein identifications had a 1% or 4% probability of being a false identification, respectively. The imported data was also searched with the X!Tandem (The Global Proteome Machine Organization), and the results from both were combined to increase protein identification confidence.

Data was imported into Cytoscape (v3.4.0) for visualization of protein-protein interactions mapped using the Protein Interaction Network Analysis (PINA) interactome database (release date 21/05/2014) ^58^ supplemented with a literature curated database of IAC proteins ^59, 60^. Three proteins could not be mapped (E9PAV3, NACA; Q9BQ48, MRPL34; E9PRG8, c11orf98) as these were not present in the PPI database.

Over-representation of gene ontology (GO) term analysis was performed using the Cytoscape plug-in ClueGO (version 2.3.3), with KEGG (Kyoto Encyclopedia of Genes and Genomes) and Reactome pathway terms. GO term grouping was used to combine related terms into groups, with a designated leading term.

### Phospho-proteomic analysis of αVβ6-dependent kinase signalling LAP-ligand stimulation assay

The TNBC BT-20 cell line was seeded in 10 cm culture dishes at 5 × 10^6^ cells per dish in complete culture medium and left to adhere overnight under standard culture conditions (37°C/8% CO2). Adherent cell monolayers were washed in TBS at RT prior to 4 hr serum starvation under standard tissue culture conditions in the presence of low serum media comprising charcoal-stripped FBS supplemented with 25 mM HEPES pH 7.0 – 7.6 (1%FBS/α-MEM/25 mM HEPES). Following serum starvation, cell monolayers were treated with human recombinant LAP (0.5 μg ml-1, L3408, Sigma Aldrich) for 10 min whilst buried on wet ice and placed on an orbital shaker in a walk-in cold room at 4°C. Monolayers were washed (x3) in pre-chilled serum-free α-MEM to remove all unbound ligand. Complete media (15% charcoal stripped FBS/α-MEM) supplemented with 25 mM HEPES was equilibrated to 37°C and added to the 30’, 15’ and 5’ time-point dishes respectively (5 ml per dish) and returned to standard culture conditions (8%CO2/37°C /humidified atmosphere) to permit ligand internalisation. The baseline control time-point was not subject to internalisation.

Ligand internalisation was quenched by flooding with TBS pre-chilled to 4°C and supplemented with phosphatase inhibitors: 1 mM sodium orthovandate (Na_3_VO_4_) and 0.5 mM sodium fluoride (NaF). A total of four biological replicates were performed across two experiments; each experiment comprised two biological replicates run in tandem.

### Cell lysis, peptide digestion and solid-phase extraction

Cells were lysed and manually harvested by scraping on wet ice in the presence of pre-chilled (4°C) phospho-proteomic lysis buffer comprising 8 M urea/20mM HEPES pH 8.0 supplemented with: 0.5 M NaF, 0.1 M Na_3_VO_4_, 1M disodium β-glycerophosphate (CH_3_H_7_Na_2_O_6_P) and 0.25 M disodium pyrophosphate (Na_2_H_2_P_2_O_7_). Lysates were sonicated at 20% intensity for 3 × 10 s prior to centrifugation at 13 000 rpm for 10 min at 4°C.

Lysis supernatants were decanted into Lo-bind microcentrifuge tubes using low-binding pipette tips to preserve protein yield. Lysates were then stored at −80°C prior to protein concentration assay in preparation for peptide desalting, titanium oxide (TiO_2_) phosphoenrichment, solid-phase extraction and interrogation of phosphopeptides by LC-MS/MS.

Lysates were thawed on wet ice and protein concentrations determined using a Pierce™ BCA Protein Concentration Assay in accordance with manufacturer’s guidelines. Samples were then normalised to a final concentration of 250 μg protein in 250 μl sample volume (1 μg μl-1) in prechilled phospho-proteomic lysis buffer.

Cysteine residues were reduced in the presence of 10 mM dithiothreitol (DTT) for 30 min in the dark at RT prior to alkylation in the presence of 40 mM iodoacetamide (IA), again for 30 min in the dark at RT. Proteins were digested in suspension using L-(tosylamido-2-phenyl) ethyl chloromethyl ketone (TPCK)-treated trypsin immobilised on agarose resin beads which had been pre-conditioned by washing and centrifugation (2000 g, 5°C, 5 min) in 20 mM HEPES pH 8.0. Samples were incubated with immobilised trypsin beads for 16 hr at 37°C with constant agitation to facilitate protein digestion.

Trypsin beads were then removed by centrifugation [2000 g, 5°C, 5 min]. The resulting peptide solutions were desalted by reversed solid-phase extraction (SPE) using OASIS® HLB (Hydrophilic-Lipophilic Balance) Extraction Cartridges rigged to a 12-port Visiprep™ SPE Vacuum Manifold to control flow rate (P=5.0 inHg ±0.5 inHg).

Columns within the cartridge were first conditioned each with 1 ml 100% acetonitrile (ACN) before equilibration with 1 ml 1% ACN/0.1% TFA (v/v) prepared in molecular grade water (mH2O; W4502, Sigma Aldrich) and washing with 500 μl 1% ACN/0.1% TFA (v/v) in mH_2_O. Peptide samples were then loaded into conditioned, equilibrated cartridge columns and purged at a low flow rate. Columns (with peptides now bound to sorbent) were washed in 1 ml 1% ACN/0.1% TFA (v/v) in mH_2_O prior to elution of desalted peptides in 250 μl 1 M glycolic acid (G8284, Sigma Aldrich) prepared in a solution of 50% ACN/5% TFA (v/v) ready for phosphopeptide enrichment.

### TiO_2_ metal oxide affinity chromatography (MOAC)

Samples were enriched for phosphopeptides by metal oxide affinity chromatography (MOAC) using titanium dioxide (TiO_2_). Desalted peptides were normalised to 500 μl with 1 M glycolic acid/80%ACN/5%TFA. Next, 125 mg dry weight of Titansphere® TiO 10 μm particles (equivalent to 12.5 mg particles per sample) was reconstituted in 250 μl 1% TFA and vortexed to ensure a homogenous suspension of Titansphere® particles. Normalised peptide samples were then incubated with 25 μl of this Titansphere® slurry (equivalent to 12.5 mg Titansphere® particles per sample) for 5 min at RT with constant agitation. Next, the enriched phosphopeptides were eluted by centrifugation [2 min, 1500 rcf] using TopTip™ micro-spin columns (TT3, Glygen Corp) previously washed with 100% ACN. Titansphere® particles were sequentially washed on a gradient accordingly: 1 M glycolic acid/80% ACN/5% TFA (x 1), 100 mM ammonium acetate (NH_4_CH_3_CO_2_)/25% ACN (x 1) and 10% ACN (x 3).

Bound phosphopeptides were eluted from Titansphere® particles by washing with 50 μl 5% NH4OH/10% ACN/mH2O per elution and centrifuging for 2 min at 1500 rcf. A total of 5 elutions were performed. Once eluted, phosphopeptide samples were immersed in dry ice to permit sublimation of NH_4_OH and desiccated overnight using a Vacufuge® Vacuum Concentrator (Eppendorf Ltd). Samples were then stored at −80°C ready for mass spectrometry.

### Nanoflow-liquid chromatography tandem mass spectrometry

Immediately prior to use, samples for analysis were reconstituted with 20 μl 50 nM enolase peptide digest (MassPREP™ Enolase Digest Phosphopeptide Mix) dissolved in 5% ACN/0.1% TFA, subjected to bath sonication (15 min, RT) and centrifugation (5 min, 5°C) to recover supernatants for LC-MS analysis.

Phosphopeptide liquid chromatography (LC) separations were performed using a Dionex UltiMate™ 3000 RSLCnano UHPLC system with an Acclaim™ PepMap™ 100 C18 RSLC Analytical Column (75 μm × 25 cm, 3 μm, 100Å) and Acclaim™ PepMap™ 100 C18 Trap Column (100 μm × 5 cm, 5 μm, 100Å). Solvents used for LC separation comprised; solvent A: 2% ACN/0.1% formic acid (FA) and solvent B: 80% ACN/0.1% FA. Sample injections of 3 μl were loaded onto the trap column at a flow rate of 8 μl min-1 over 5 min. After loading, samples were eluted along an 85 min gradient from 6.3% to 43.8% solvent A prior to column cleaning in 90% solvent B for 10 min and equilibration with 6.3% solvent A for 10 min.

All analyses were performed on the Thermo Scientific LTQ Orbitrap™-Velos™ Hybrid FT spectrometer. The instrument was operated in data-dependent acquisition (DDA) mode, whereby a full MS1 survey scan (m/z 350 – 1500) was completed at a resolution of 30 000 FWHM (m/z 400) and ions were analysed in the Orbitrap™. The top 7 most intense multiply charged precursor ions detected in the MS1 survey scan were automatically mass-selected for fragmentation by collision-induced dissociation (CID) with multi-stage activation enabled and analysed in the LTZ-Velos™ linear ion trap (m/z 190 – 2000). Dynamic exclusion was enabled to prevent repeat analysis of identical precursor ions within 60 secs.

### Identification and quantification of phosphopeptides

Mascot Daemon and Distiller software (v2.3.0.0 and v2.4.2.0 respectively, both Matrix Science) were used to convert exported LTQ Orbitrap™-Velos™. raw files into. mgf files for peak list searches against the UniProtKB/SwissProt human proteome database (UniProtConsortium 2017). Data were searched according to the following criteria: ±10 ppm precursor and ±600 mmu fragment ion m/z tolerances; digestion enzyme = trypsin (2 missed cleavages tolerated); fixed modification: carbamidomethyl (C); variable modifications: oxidation (M), phospho (ST), phospho (Y) and gln -> pyro-glu (glutamine -> pyroglutamate) (Q) at N-terminus.

Mascot search engine results for phosphopeptide identification were collated using Perl script (Perl® 5, Perl.org) and Post Analysis Data Acquisition v1.1 (PAnDA) software^61^. Data were curated algorithmically to include only unique phosphopeptide ions with a q-value ≤0.05, calculated by comparison to searches against a randomised database. All phosphopeptides assigned a Mascot delta score ≥10 were reported as the specific phosphorylation site.

Phosphopeptides were quantified using PEak Statistical CALculator (PESCAL)^62^ to generate extracted ion chromatograms for the first three isotopes of each phosphopeptide ion within the database (±7 ppm m/z tolerance, ±1.5 min retention time tolerance, isotope correlation >0.8) enabling calculation of peak height for each constructed extracted ion chromatogram.

Using the R statistical programming environment (R 3.2.5, The R Foundation), the peak heights for each phosphopeptide ion were then log2-transformed, quantile normalised and fitted to a linear model where difference in magnitude and statistical significance between time-points was calculated using Bayes shrinkage of standard deviations^63^. The p-values generated were then subject to Benjamini-Hochberg post-hoc analyses to correct for multiple testing. Kinase Substrate Enrichment Analysis (KSEA) was performed on the dataset as previously described^25^, in order to infer phospho-proteomic network activity and plasticity.

### Indirect Immunofluorescence

Cells were fixed with 4% (w/v) paraformaldehyde (PBS (-), pH 6.9) for 20 minutes at room temperature, then washed three times in PBS (-). Cells were then permeabilized for 3 – 4 minutes with 0.1% (v/v) Triton X-100 at room temperature followed by three 0.1/0.1 buffer washes. Primary antibody incubations were for 45 minutes at room temperature, followed by three washes in 0.1/0.1 buffer. Samples were incubated with secondary antibody, with or without phalloidin (1:400 in 0.1/0.1 buffer), for 45 minutes at room temperature protected from light. Samples were then washed twice in PBS (-) and once in Milli-Q water, before mounting with Prolong Gold anti-fade mountant (Molecular Probes Invitrogen) on glass Superfrost® Plus glass slides (Thermo Scientific).

Samples were imaged using a Zeiss 3i Marianas spinning disk confocal system using a 63x/1.4 oil objective. Downstream image processing was performed using Image J FIJI.

### Inhibition of Protein Degradation

For LAMP2 co-localisation assays, cells were pre-incubating with folimycin and epoxomicin to inhibit lysosomal and proteasomal degradation, respectively ^64^. Folimycin (Calbiochem) was used as a lysosomal inhibitor, as it is a potent and selective inhibitor of the vacuolar-type H+-ATPase that prevents acidification of the lysosome. Epoxomicin (Calbiochem) was used as a proteasomal inhibitor, as it is a potent and selective inhibitor of the peptide hydrolysing activities of the proteasome. Serum starved cells were incubated in 100 nM concentrations of both folimycin and epoxomicin for 6 hours, before EGF or LAP stimulation ^64^. Inhibitor stocks were reconstituted in DMSO (1 mg/ml) therefore control conditions included a DMSO vehicle control.

### Surface Receptor Internalisation Assay

Cells were seeded at a 60% density in 10 cm dishes (three per condition) for 8 hours, before serum starvation overnight, by washing three times in PBS (-) then incubating in DMEM/25mM HEPES solution. High binding half area 96 well ELISA (Enzyme Linked Immunosorbent Assay) immunoassay plates (Corning) were coated overnight at 4°C on a rocker with 5 μg/ml primary antibody (620W7 anti-integrin αVβ6, rat polyclonal) diluted in 0.05 M NaCO_3_ pH 9.6. The 96 well plates were washed four times in PBST, then blocked in 5% BSA at room temperature for a minimum of 1 hour. All wash steps with cells during the assay were done on ice with ice cold buffers, unless otherwise stated.

Cells were washed twice in Krebs, before labelling with EZ-Link™ Sulfo-NHS-SS-Biotin (Thermo Scientific) (220 μM in PBS (-)) at 4°C on a gentle rocker (7 RPM). Unbound biotin was removed by washing three times in Krebs. ‘Total’ positive control cell plates were returned to 4°C to show maximal surface biotinylated receptor signal. ‘Blank’ negative control plates were also returned to 4°C to show the efficiency of surface biotin removal without internalisation. Equilibrated warm medium alone or with EGF (10 ng/ml) was added to plates, before immediate transferral to 37°C to allow surface receptor internalisation for either 4, 7, 15 or 30 minutes.

After internalisation plates were immediately returned to ice and washed twice with PBS (-) and once with pH 8.6 buffer (100 mM NaCl, 50 mM Tris pH 7.5, pH 8.6 at 4°C). pH 8.6 buffer supplemented with 22.84 mM Sodium 2-mercaptoethanesulfonate (MesNa) (Fluka Analytical) and 0.22 mM NaOH was then added to the blank and internalised plates, before incubation for 30 minutes at 4°C on a gentle rocker. The incubation with the reducing agent MesNa removes surface biotin by cleaving the reducible disulphide bond in the biotin reagent. Biotin bound to internalised surface receptors is unaffected, as MesNa is cell-impermeant^65^.

Plates are then washed twice in PBS (-) then lysed in lysis buffer (200 mM NaCl, 75 mM Tris, 7.5 mM EDTA, 7.5 mM EGTA, 1.5% (v/v) Triton X-100, 0.75% (v/v) Igepal CA-630, 15 mM NaF, 1.5 mM Na_3_VO_4_, 50 µg/µl leupeptin, 50 µg/µl apoprotein, 1 mM AEBSF). Lysates were spun at 13,000 × g for 10 minutes at 4°C. The blocking buffer was removed from the 96 well plate which was washed four times in PBST, before the lysate supernatant was added into each corresponding well. Plates were incubated with the lysate overnight at 4°C.

Unbound material was removed with four PBST washes. Wells were then incubated with streptavidin-conjugated horseradish peroxidase (1:500) in PBST containing 1% BSA, for 1 hour at room temperature. The plate was then washed again four times in PBST, before incubation with an ABTS substrate solution (ABTS buffer: 0.1 M sodium acetate, 0.05 NaH_2_PO_4_ pH 5, with 2 mM ABTS, 2.5 mM H_2_O_2_). The resultant colourimetric change was measured at 405 nm absorbance on a Thermo Labsystem Multiskan Spectrum plate reader. Readings were taken across multiple time points ranging from 30 minutes to 24 hours, depending on the rate of development.

### Proximity Ligation Assay

Duolink In Situ Fluorescence Detection Reagents Orange (Olink® Bioscience), a proximity ligation assay, was utilized to detect and visualize protein interactions between αvβ6 and EGFR. Briefly, cells were seeded onto coverslips and formalin-fixed 24h later. Non-specific binding was blocked using 5% goat serum in 1% PBS 0.1% Tween-20, followed by incubation with primary antibodies to β6 and EGFR (10D5 and D38B1 (NEB), respectively) overnight at 4°C. After washing, cells were incubated for 1h 37°C with PLA probes (secondary antibodies to mouse (for 10D5) and rabbit (for EGFR) conjugated with a PLA probe). Cells were incubated with Ligation-ligase solution (40 min, 37°C) and any interacting oligonucleotides amplified with Amplification-Polymerase solution (100 min, 37°C). Cells were washed & mounted onto slides using mounting media containing DAPI (from Duolink kit). Molecular proximity was visualized using fluorescence confocal microscopy.

### Traction Force Microscopy

Polyacrylamide hydrogels embedded with FluoSpheres® carboxylate-modified 0.2 μm fluorescent (505/515) microspheres were synthesised as described previously^32^. Hydrogels were treated with sulfo-SANPAH under UV light (365nm), then coated with LAP (0.5μg/ml) overnight at 4°C. 9×10^4^ cells were seeded per hydrogel and incubated overnight in serum-free media at 37°C with 5% CO_2_. Point visiting was used to visualise live cells and beads on a Zeiss 3i Marianas spinning disk confocal system using a 63x/1.4 oil objective. Z-stack images were taken for each cell and underlying microspheres within the hydrogel 1) prior to stimulation with EGF (10ng/ml), 2) after 30 minutes EGF stimulation, and 3) following lysis of cells with 1% SDS. Images were analysed in ImageJ. Analysis consisted of processing images via alignment, registration and cropping, followed by particle image velocimetry (PIV) and Fourier Transform traction cytometry (FTTC) analysis^32, 66^.

### RhoA activity Assay - G-LISA

MDA-MB-468 cells were plated on FN-coated 10cm dishes in DMEM for 2-4 Hrs prior to treatment with 10 μg/ml Rat IgG or anti-αVβ6 blocking antibody (53A2). Treatments were staggered to ensure all time-points (0, 30, 60 and 120 mins) were lysed simultaneously. Following lysis, in G-LISA buffer (cytoskeleton) lysates were snap-frozen in N_2(l)_. RhoA activity was assessed using absorbance-based G-LISA RhoA Activation Assay Kits (Cytoskeleton; #BK124) according to manufacturer’s instructions.

### Plating cells in 3D collagen/FN/LAP matrices

Collagen gel solutions with a final concentration of 1.7 mg/m or 3 mg/m Rat Tail Collagen I were prepared. Gels were supplemented with Dulbecco’s Modified Eagle Medium (1x;from 10x stock) and buffered with Sodium Bicarbonate (3.7 g/l) before addition of Bovine Fibronectin (25 ug/ml) and Human Transforming Growth Factor-β1 Latency Associated Peptide (0.5 μg/ml). For each condition 1 × 10^5^ cells were suspended in 200 μl of Collagen mixture and seeded onto cell imaging dishes (Stratlab Ltd., ref: 450-AK-005). Gels were then allowed to polymerise at 37°C for 1 hr before the addition of 500 μL of growth medium containing 10% serum. Following overnight incubation, gels were washed thouroughly with DMEM and cells serum-starved for 2.5 hrs prior to 1 hr treatment with 10 μg/ml Rat IgG or anti-αVβ6 blocking antibodies, or 3 hrs prior to 30 min treatment with vehicle control or EGF (10 ng/ml).

### Statistical Analysis of cell biological data

Pair-wise comparisons of continuous data were tested using Student’s two-tailed t-test for parametric data, or Rank sums test for non-parametric data where the equal variance test (Brown-Forsythe) failed. Comparisons between more than two groups were tested using the analysis of variance (ANOVA) with the appropriate *post-hoc* test for multiple test correction. Statistical tests were performed using either SigmaPlot 13.0 statistical software, or within the ‘Quantitative Analysis’ section of Scaffold 4.8 (Proteome Software).

### Transwell invasion assays

5×10^4^ cells were seeded per well post-treatment into 6.5mm diameter, 8μm pore-sized Transwells® (Corning BV) coated with 70μl BD Matrigel Basement Membrane matrix (Matrigel):media (1:2 ratio). Cells which invaded through Matrigel were counted after 72h using a CASY counter (Scharfe Systems, Germany).

## ACKNOWLEDGEMENTS

This work was supported by a Breast Cancer Campaign grant BCC 2008MayPR32 (KMM), Wellcome Trust PhD studentships 102379/Z/13/Z (JRT) & 105350/Z/14/Z (DN), a Cancer Research UK PhD studentship (CS), Breast Cancer Now Grant 2015MayPR507 (HM) and North West Cancer Research fellowship (DH)

The mass spectrometers and microscopes used in this study were purchased with grants from Cancer Research UK, Wellcome Trust and University of Liverpool (Institute of Translational Medicine) Strategic Fund.

## SUPPLEMENTARY FIGURES

**Figure S1:**
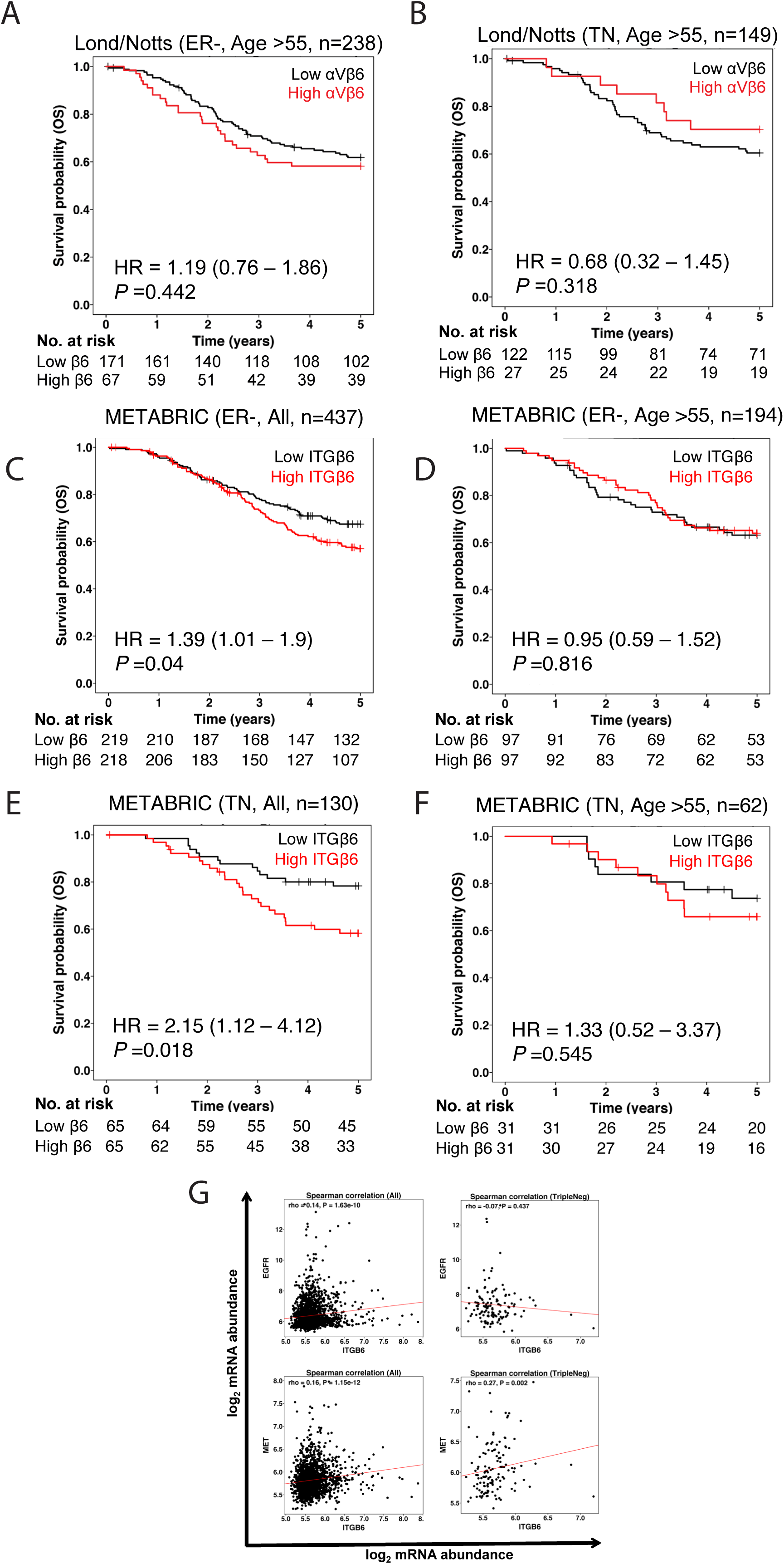
Expression of ITGB6 and EGFR genes in breast cancer patients by subtype. Kaplan-Meier curves by integrin αVβ6 expression status. Tick marks indicate patients who were censored. All *P* values refer to log-rank tests. **A & B)** Cancerous breast cancer tissue sections were immunohistochemically stained for integrin αVβ6 using 6.2G2 antibody. Overall survival in both London and Nottingham combined cohorts (Lond/Notts) of breast cancer patients by integrin αVβ6 status (high αVβ6 expression: red; low αVβ6 expression: black). **A**) Overall survival of ER-negative patients (ER-) >55 years of age by integrin αVβ6 status. **B)** Overall survival of triple negative (TN) breast cancer patients >55 years of age by integrin αVβ6 status. **C-F)** Overall survival in METABRIC by ITGB6 gene status (high expression in red, low in black). **C)** The *P* value for all ER-negative (ER-) patients with high ITGB6 gene expression versus low expression tumours is *P*=0·04 for all age groups. **D)** Overall survival of ER-patients >55 years of age. **E)** The *P* value for all triple negative (TN) breast cancer patients with high ITGB6 gene expression versus low expression is *P*=0·018. (**F**) Overall survival of TN patients >55 years of age. **G)** *ITGB6*, and *EGFR* gene expression in breast cancer patients by subtype. Spearman correlation of ITGB6 gene status in the METABRIC dataset. *EGFR* mRNA profiles demonstrated a modest trend of significant positive correlation with *ITGB6*.

**Figure S2:**
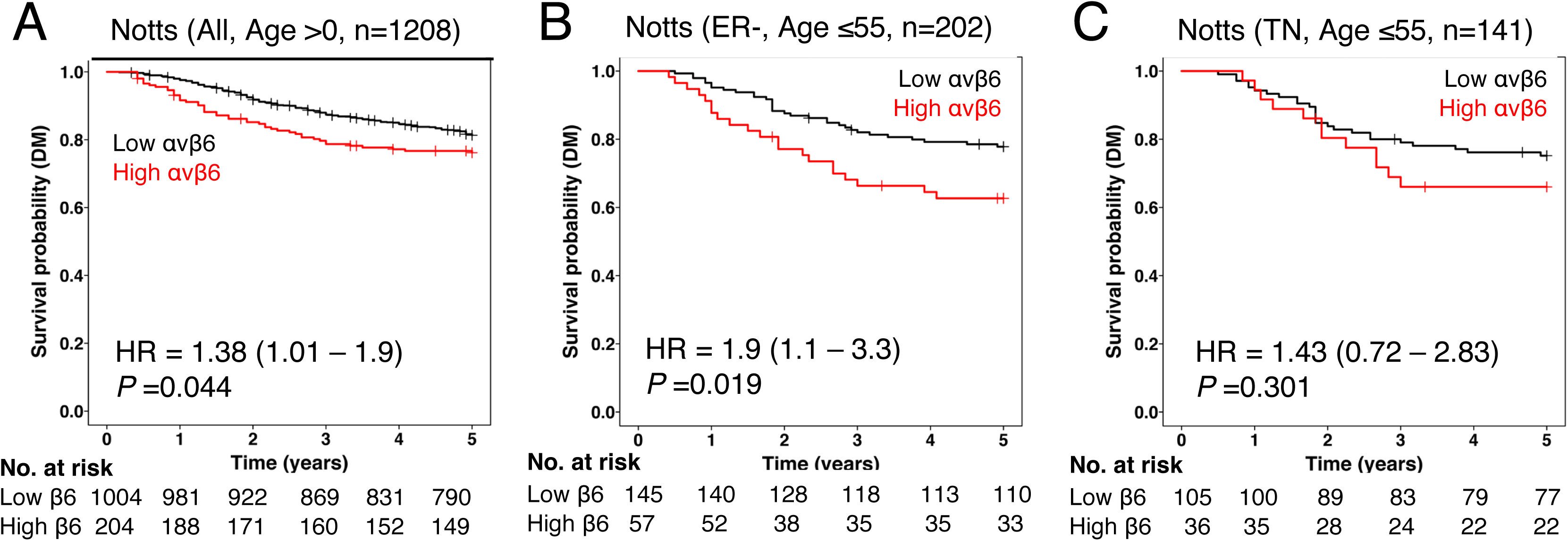
Expression of integrin αVβ6 and overall survival in breast cancer patients with distant metastasis. Kaplan-Meier survival curves by integrin αVβ6 expression status in patients with distant metastasis. Tick marks indicate patients who were censored. All *P* values refer to log-rank tests. Cancerous breast cancer tissue sections were immunohistochemically stained for integrin αVβ6 using 6.2G2 antibody (Biogen Idec). Overall survival in the Nottingham (Notts) cohort of breast cancer patients by integrin αVβ6 status (high expression in red, low in black. **A**). Overall survival of the cohort. **B)** Overall survival of ER-negative (ER-) patients with distant metastasis ≤55 years of age. The *P* value for ER-negative patients with distant metastasis ≤55 years with high tumor integrin αVβ6 expression versus low αVβ6 expression is 0·019, with an HR=1·9 (95%CI 1·1-3·3) for those with higher expression. **C)** Overall survival of triple-negative (TN) breast cancer patients from the Nottingham patient cohort ≤55 years of age by integrin αVβ6 status.

**Figure S3:**
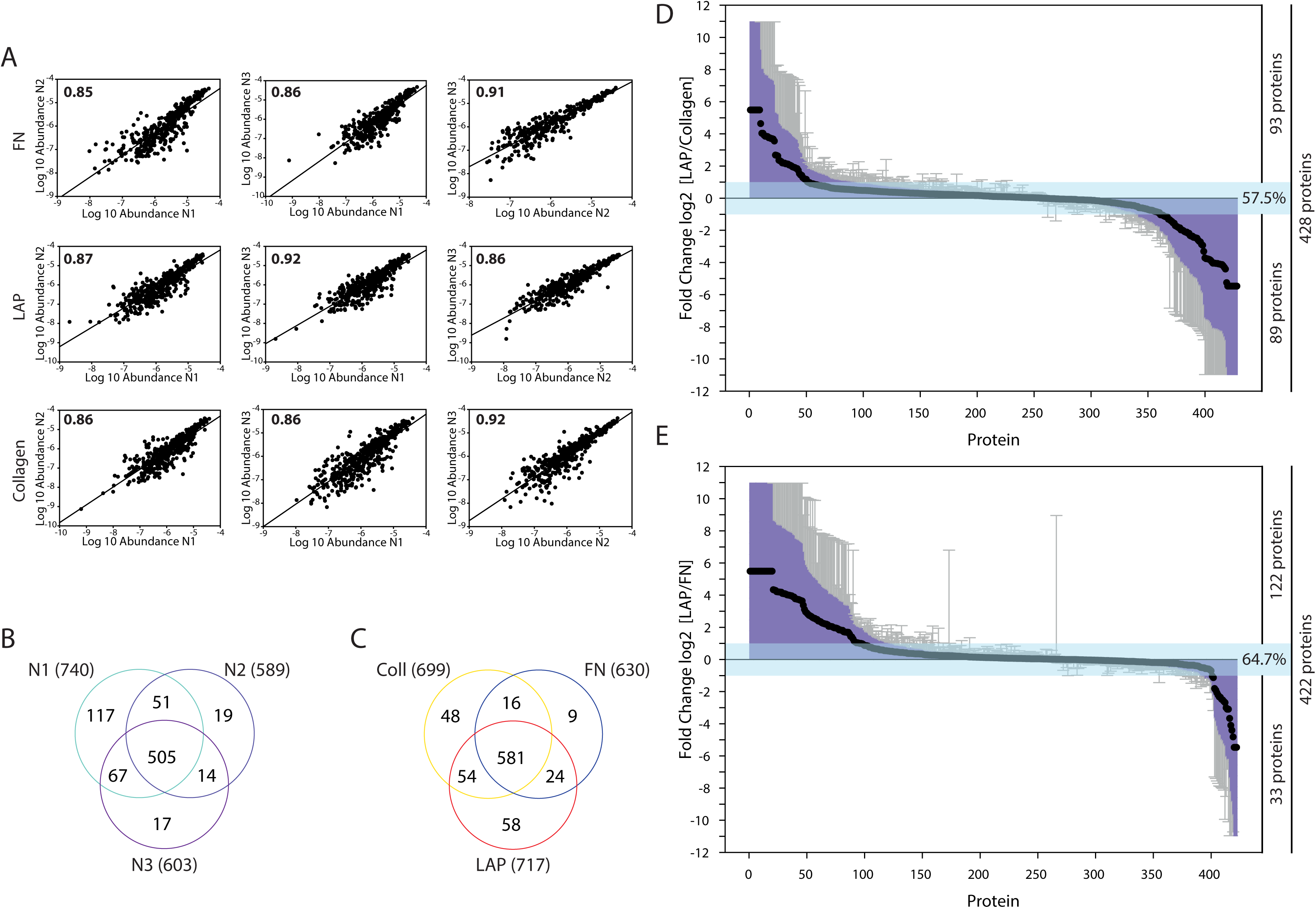
Quality assurance of proteomic datasets. **A)** Scatter plots with regression lines. Pairwise comparison of log_10_ transformed protein abundance normalized spectral abundance factor (NSAF) values between replicates for fibronectin, LAP and collagen ligand samples. Pearson correlation coefficient value is indicated in the top left of each graph. **B & C)** Venn Diagrams demonstrating the degree of shared and unique proteins between replicates (B) and ligands (C). Replicates indicate the total number of proteins pooled from all three ligands from each biological replicate. Ligands indicate the total number of proteins isolated from adhesions on each ligand from all three biological replicates. Total proteins identified = 790. **D & E)** Fold change of proteins between ligand conditions. Low abundance (fewer than 5 peptides and/or unique to one N) and unreviewed UniProt proteins were removed from the dataset, yielding a total of 428 proteins. Graphs show ratios of normalised intensity values for log_2_ (D) [LAP/Collagen] and (E) [LAP Vs FN], mean ± SEM, N = 3 for each protein. Light blue shading corresponds to ≤ two-fold change. (D) LAP Vs Collagen, 428 proteins in total, 57.5% of dataset (93 and 89 proteins increased and decreased, respectively). (E) LAP Vs FN, 422 proteins in total, 64.7% of dataset (122 and 33 proteins increased and decreased, respectively).

**Figure S4:**
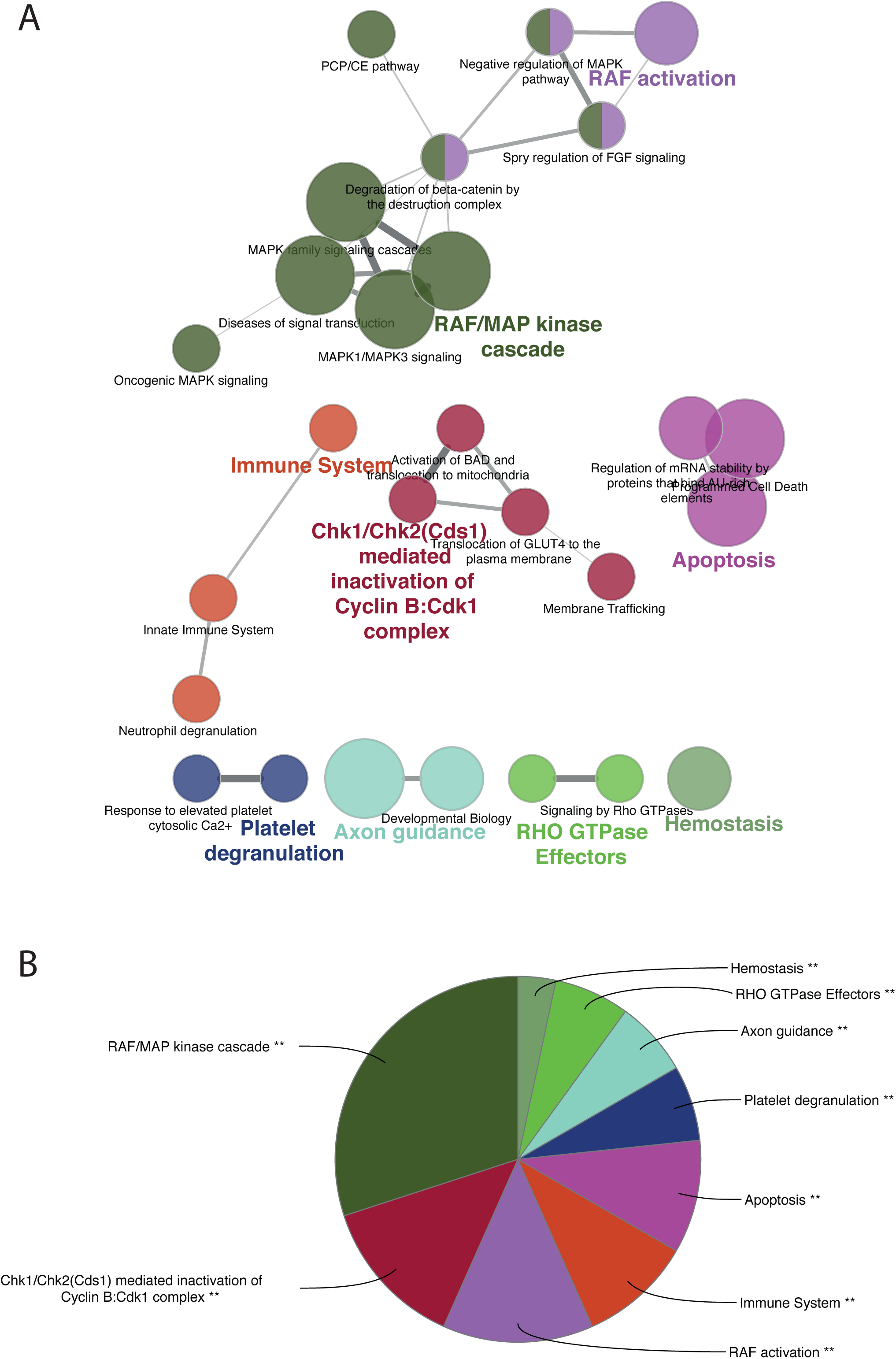
Reactome analysis of αVβ6-dependent adhesion complexes. ClueGO Reactome Pathway term hierarchical clustering identifies RAF/MAP kinase cascade in αVβ6 IACs. **A)** ClueGO hierarchical layout of represented Reactome pathway terms (Pathway level 3-8). Node colour corresponds to grouping, with the lead term in corresponding coloured text. Nodes with split colours belong to multiple groups. Nodes represent individual Reactome pathway terms. **B)** Pie-chart organised by the % of genes per term. Analysis parameters: minimum 1 gene per cluster, GO term/pathway network connectivity (Kappa score) =1; Statistical test Enrichment/Depletion (Two-sided hypergeometric test), Bonferroni p-value correction, p-value threshold <0.05 applied; GO term grouping based on kappa score, 50% of genes for group merge, 50% terms for group merge. Leading group term based on %gene/term.

**Figure S5:**
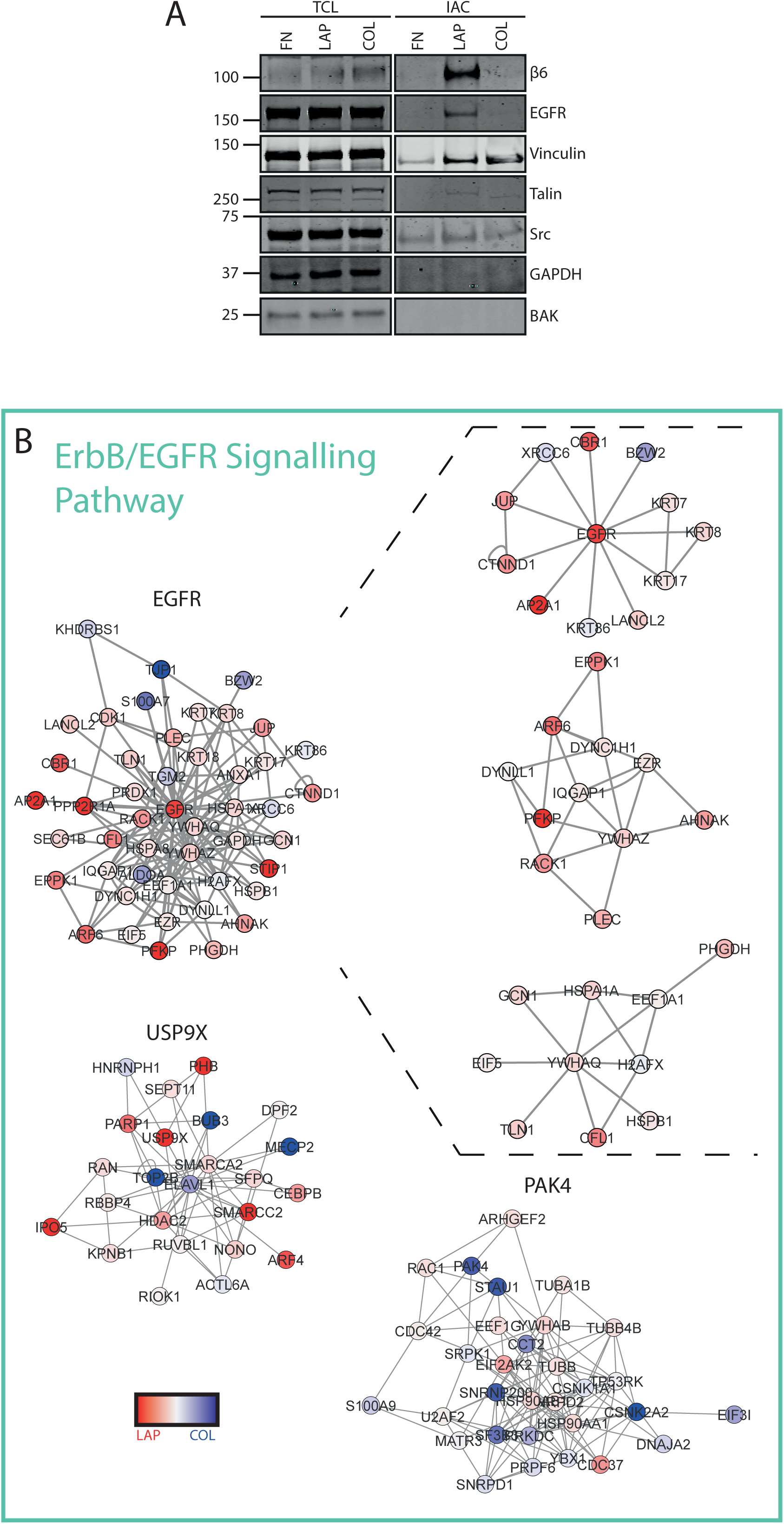
ErbB/EGFR signalling pathway term subnetwork clusters in αVβ6-dependent IACs. **A)** EGFR is detected in αVβ6-dependent IACs on LAP. Immunoblotting total cell lysate (TCL) and isolated integrin-associated complexes (IAC) for the EGFR, adhesion complex components (integrin β6, vinculin, talin & Src), and negative control proteins (GAPDH and BAK).**B)** Key clusters of proteins identified in αVβ6-dependent IACs on LAP associated with ErbB/EGFR signalling pathway KEGG term and their one-hop interactors. Dashed lines from the EGFR primary cluster identify sub-clusters derived from the EGFR primary cluster. Primary clusters are named according to key proteins. Inter-cluster interactions are not shown. Nodes represent proteins, and edges are known interactions. Node colour red to blue gradient = log_2_ fold enrichment LAP/collagen.

**Figure S6:**
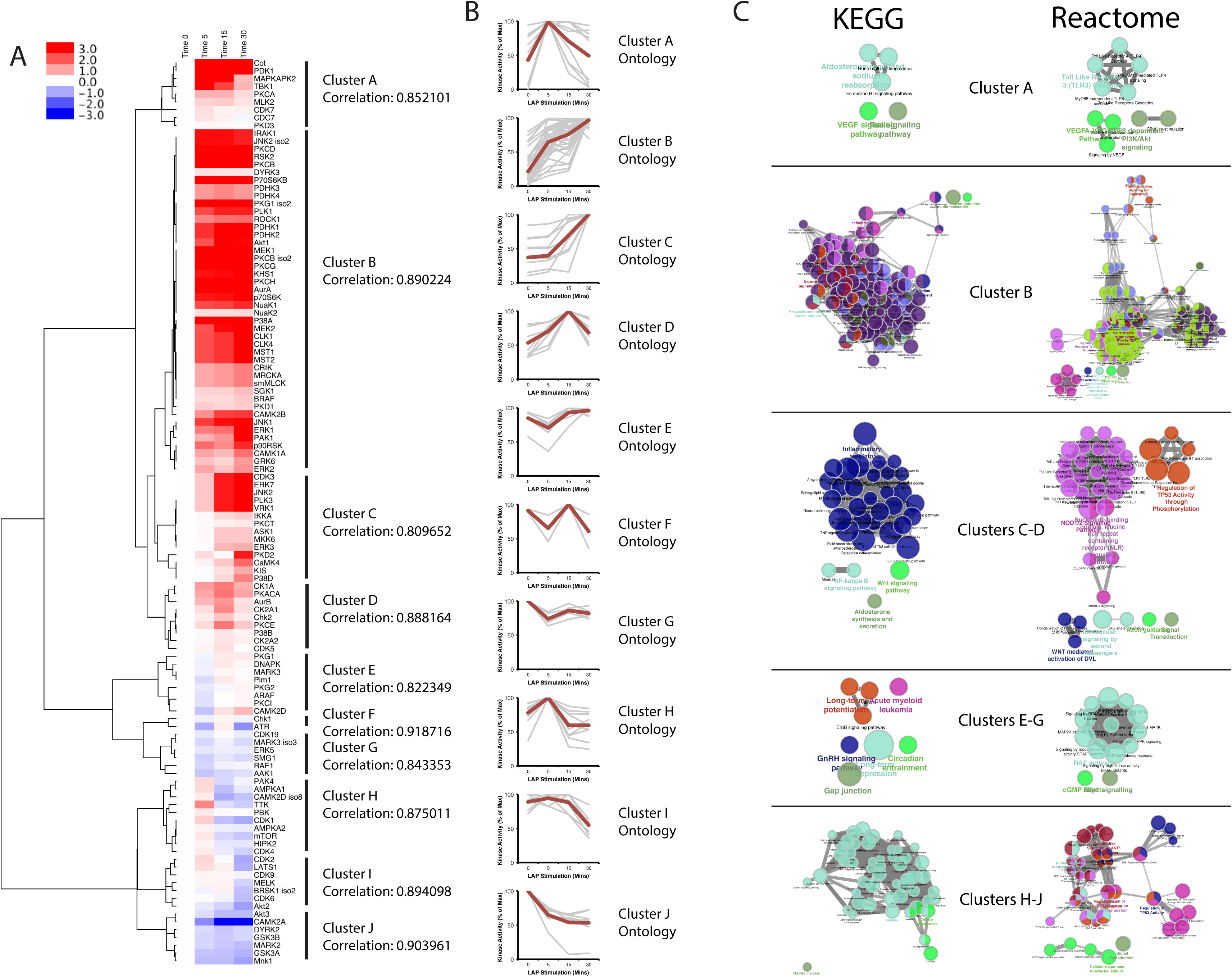
KSEA Clustering - KEGG & Reactome. Kinase activation following ligand-dependent αVβ6 stimulation reveals αVβ6-EGFR crosstalk. Kinase Substrate Enrichment Analysis (KSEA) in BT-20 TNBC cells following stimulation of αVβ6 with soluble latency-associated peptide (LAP) (Time points: 0, 5, 15 & 30 mins), to infer kinase network plasticity during integrin α?β6 LAP-engagement **A)** Hierarchical clustering of differentially activated/inactivated predicted kinases. Unsupervised hierarchical clustering of individual kinases within the KSEA repository identified as significantly regulated (activated or inactivated) following LAP stimulation. Clustering based on activation/inactivation profiles. Cluster threshold: Spearman’s rank correlation coefficient ρ≥0.8 Summary heatmap of individual kinases in each cluster. Colour scale: Mean average kinase activity (% of maximal activity) of kinases grouped within each cluster according. Red to Blue gradient: Active to Inactivation scale **B)**Profile plots for each cluster with the mean temporal profile for each cluster indicated by a red line. **C)**ClueGo heirarchical layout of KEGG & Reactome pathway terms (Pathway level 3-8) for individual clusters or groups of clusters (Cluster A, Cluster B, Clusters C-D, Clusters E-G, Clusters H-J

**Figure S7:**
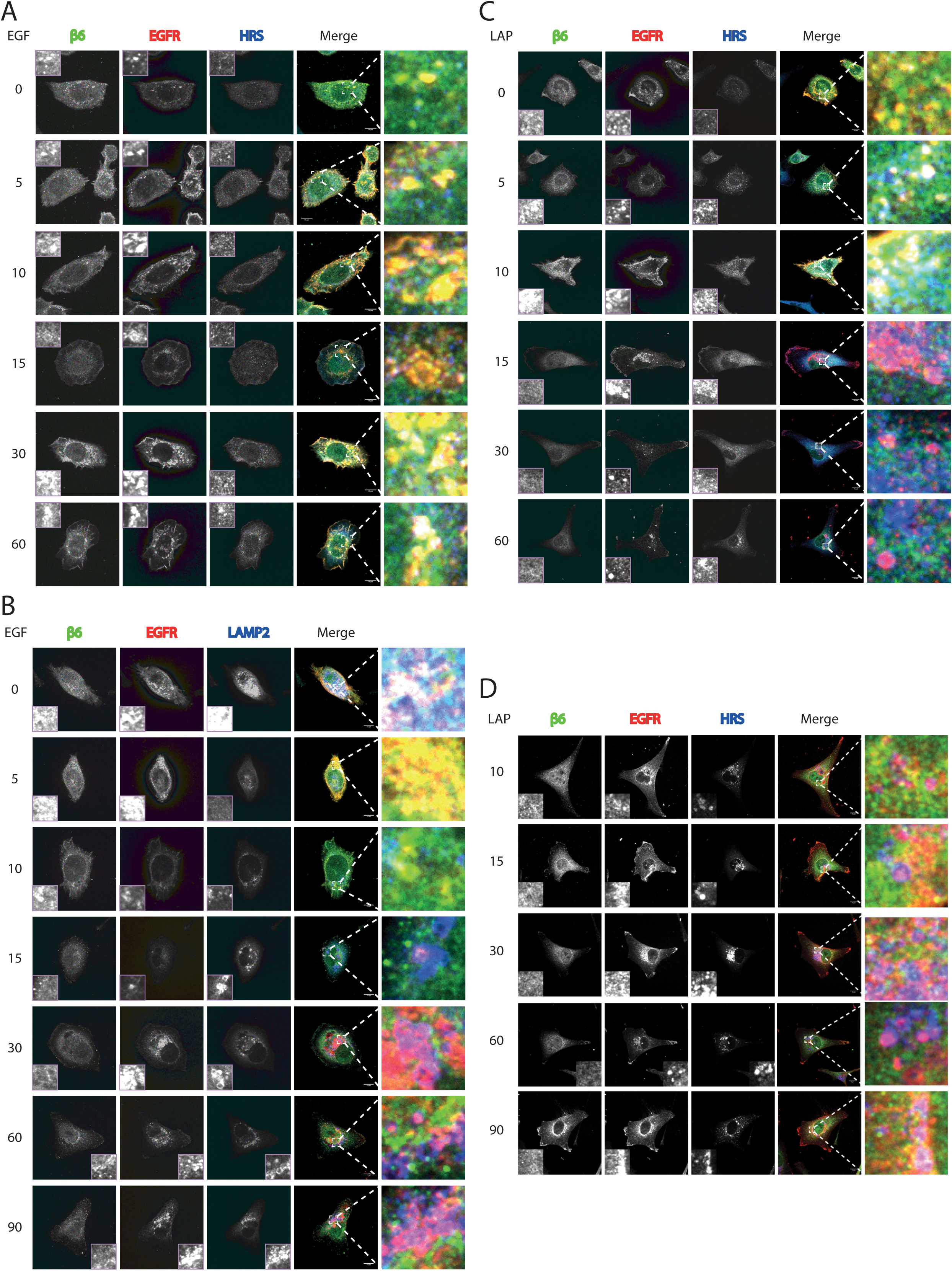
αVβ6-EGFR crosstalk regulates reciprocal receptor trafficking mechanisms. **A)** EGF stimulation induces alisation of β6 and EGFR with HRS. MDA-MB-468 cells, co-stained for β6, EGFR (R-1) and HRS. Single z slice shown for a juxtamembrane section of the cell. EGF stimulation shown in minutes. Scale bar = 10 μm. **B)** EGF stimulation induces alisation of EGFR with LAMP2.MDA-MB-468 cells, co-stained for β6, EGFR (D38B1) and LAMP2. Cells were pre-incubated with the lysosomal and proteasomal inhibitors folimycin and epoxomicin, respectively. Single z-slice shown for a juxtamembrane section of the cell. EGF stimulation shown in minutes. EGFR is shown at 100 - 19000 for 0 - 10 minutes and 100 - 2600 for 15 - 90 minutes, intensity arbitrary units. Scale bar=10 μm**C)** LAP stimulation induces alisation of β6 and EGFR with HRS. MDA-MB-468 cells, co-stained for β6, EGFR (D38B1) and HRS. Single z slice shown for a juxtamembrane section of the cell. LAP stimulation is shown in minutes. for 15, 30 and 60 minutes, intensity arbitrary units. Scale bar = 10 μm **D)** LAP stimulation induces alisation of EGFR with LAMP2.MDA-MB-468 cells, co-stained for β6, EGFR (D38B1) and LAMP2. Cells were pre-incubated with the lysosomal and proteasomal inhibitors folimycin and epoxomicin, respectively. Single z slice shown for a juxtamembrane section of the cell. LAP is stimulation shown in minutes‥ Scale bar = 10 μm

**Figure S8:**
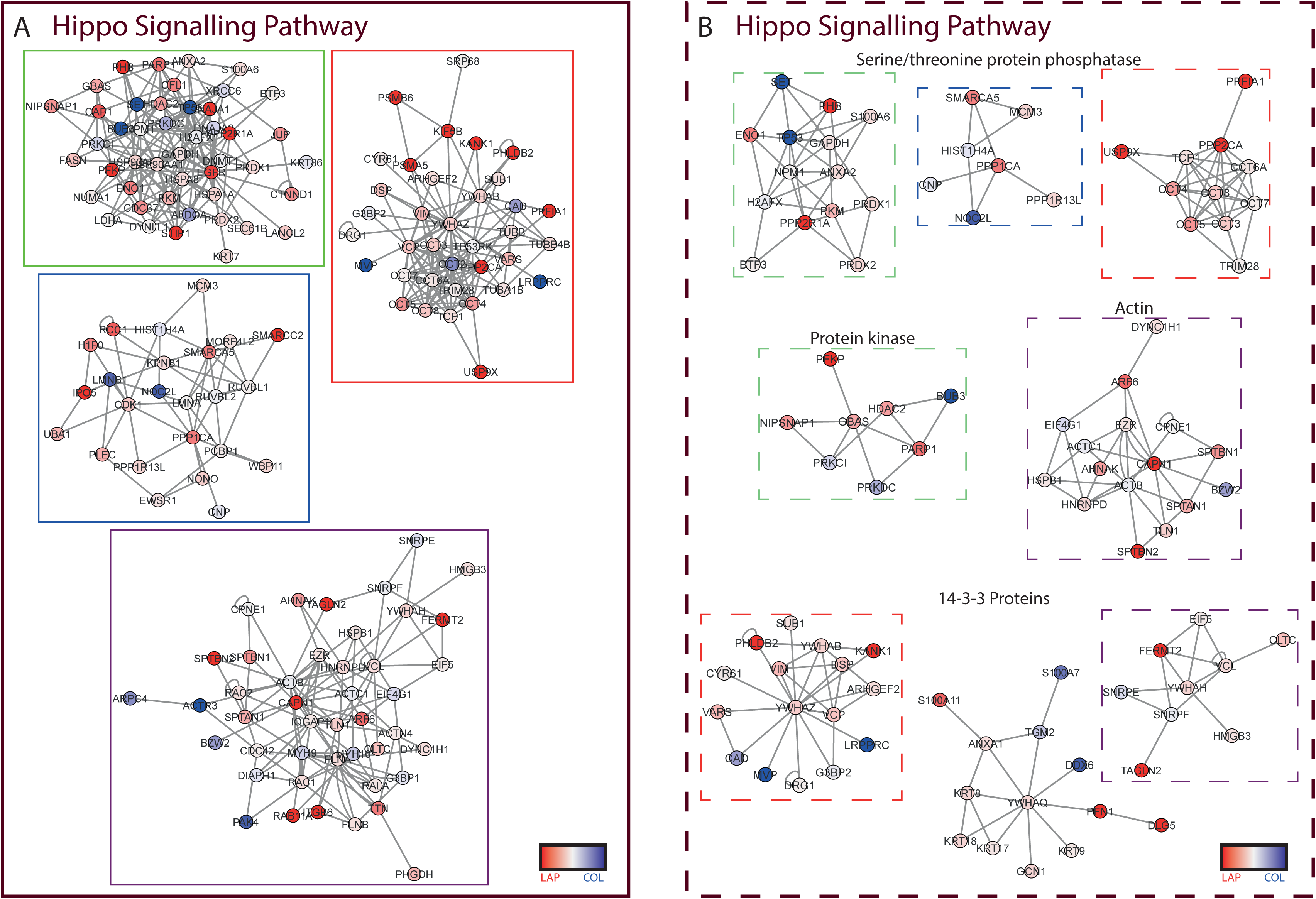
Hippo signalling pathway term subnetwork clusters in αVβ6-dependent IACs. Key clusters of proteins identified in αVβ6-dependent IACs on LAP associated with hippo signalling pathway KEGG term and their one-hop interactors. **A)** Hippo signalling pathway term primary subnetwork clusters. **B)** Hippo signalling pathway term sub-clusters derived from primary clusters. **A & B)** Clusters are colour matched: Continuous outline for the primary cluster **(A)**; Dashed outline for sub-clusters derived from primary clusters **(B)**. Clusters named according to key proteins. Inter-cluster interactions are not shown. Nodes represent proteins, and edges are known interactions. Node colour red to blue gradient = log_2_ fold enrichment LAP/collagen.

## Abbreviations

EGFR: Epidermal Growth Factor Receptor
ER: Oestrogen Receptor
FN: Fibronectin
HER2: human epidermal growth factor receptor 2
IAC: Integrin-Associated Complex
LAP: Latency-Associated Peptide
MAPK: Mitogen activated protein kinase
OS: Overall Survival
PR: Progesterone Receptor
RTK: Receptor Tyrosine Kinase
TNBC: Triple Negative Breast Cancer
YAP: Yes-Associated Protein-1
TAZ: Tafazzin

## REFERENCES

1. Butcher, D.T., Alliston, T. & Weaver, V.M. A tense situation: forcing tumour progression. Nature reviews. Cancer 9, 108–122 (2009).

2. Frantz, C., Stewart, K.M. & Weaver, V.M. The extracellular matrix at a glance. Journal of cell science 123, 4195–4200 (2010).

3. Cox, T.R., Gartland, A. & Erler, J.T. Lysyl Oxidase, a Targetable Secreted Molecule Involved in Cancer Metastasis. Cancer Res 76, 188–192 (2016).

4. Chin, V.T. et al. Rho-associated kinase signalling and the cancer microenvironment: novel biological implications and therapeutic opportunities. Expert Rev Mol Med 17, e17 (2015).

5. Pajic, M. et al. The dynamics of Rho GTPase signaling and implications for targeting cancer and the tumor microenvironment. Small GTPases 6, 123–133 (2015).

6. Humphries, J.D., Paul, N.R., Humphries, M.J. & Morgan, M.R. Emerging properties of adhesion complexes: what are they and what do they do? Trends Cell Biol 25, 388–397 (2015).

7. Desgrosellier, J.S. & Cheresh, D.A. Integrins in cancer: biological implications and therapeutic opportunities. Nat Rev Cancer 10, 9–22 (2010).

8. Moore, K.M. et al. Therapeutic targeting of integrin alphavbeta6 in breast cancer. J Natl Cancer Inst 106 (2014).

9. Bates, R.C. et al. Transcriptional activation of integrin beta6 during the epithelial-mesenchymal transition defines a novel prognostic indicator of aggressive colon carcinoma. J Clin Invest 115, 339–347 (2005).

10. Hazelbag, S. et al. Overexpression of the alpha v beta 6 integrin in cervical squamous cell carcinoma is a prognostic factor for decreased survival. J Pathol 212, 316–324 (2007).

11. Elayadi, A.N. et al. A peptide selected by biopanning identifies the integrin alphavbeta6 as a prognostic biomarker for nonsmall cell lung cancer. Cancer research 67, 5889–5895 (2007).

12. Moore, K.M. et al. Therapeutic targeting of integrin avß6 in breast cancer. J Natl Cancer Inst 106 (2014).

13. Allen, M.D., Marshall, J.F. & Jones, J.L. alphavbeta6 Expression in myoepithelial cells: a novel marker for predicting DCIS progression with therapeutic potential. Cancer Res 74, 5942–5947 (2014).

14. Morgan, M.R. et al. Psoriasin (S100A7) associates with integrin ß6 subunit and is required for avß6-dependent carcinoma cell invasion. Oncogene 30, 1422–1435 (2011).

15. Elosegui-Artola, A. et al. Rigidity sensing and adaptation through regulation of integrin types. Nat Mater 13, 631–637 (2014).

16. Breuss, J.M. et al. Expression of the beta 6 integrin subunit in development, neoplasia and tissue repair suggests a role in epithelial remodeling. J Cell Sci 108 (Pt 6), 2241–2251 (1995).

17. Howlader N, N.A., Krapcho M, Miller D, Bishop K, Kosary CL, Yu M, Ruhl J, Tatalovich Z, … Vol. https://seer.cancer.gov/csr/1975_2015/ (

18. Streuli, C.H. & Akhtar, N. Signal co-operation between integrins and other receptor systems. Biochem J 418, 491–506 (2009).

19. Ivaska, J. & Heino, J. Cooperation between integrins and growth factor receptors in signaling and endocytosis. Annu Rev Cell Dev Biol 27, 291–320 (2011).

20. Bleyer, A. et al. The distinctive biology of cancer in adolescents and young adults. Nature Reviews Cancer 8, 288–298 (2008).

21. Gnerlich, J.L. et al. Elevated Breast Cancer Mortality in Women Younger than Age 40 Years Compared with Older Women Is Attributed to Poorer Survival in Early-Stage Disease. J Am Coll Surgeons 208, 341–347 (2009).

22. Kurian, A.W., Fish, K., Shema, S.J. & Clarke, C.A. Lifetime risks of specific breast cancer subtypes among women in four racial/ethnic groups. Breast Cancer Research 12 (2010).

23. Horton, E.R. et al. Definition of a consensus integrin adhesome and its dynamics during adhesion complex assembly and disassembly. Nat Cell Biol 17, 1577–1587 (2015).

24. Casado, P. et al. Phosphoproteomics data classify hematological cancer cell lines according to tumor type and sensitivity to kinase inhibitors. Genome Biol 14, R37 (2013).

25. Casado, P. et al. Kinase-substrate enrichment analysis provides insights into the heterogeneity of signaling pathway activation in leukemia cells. Sci Signal 6, rs6 (2013).

26. Sorkin, A. & Goh, L.K. Endocytosis and intracellular trafficking of ErbBs. Exp Cell Res 315, 683–696 (2009).

27. Raiborg, C. & Stenmark, H. The ESCRT machinery in endosomal sorting of ubiquitylated membrane proteins. Nature 458, 445–452 (2009).

28. Wenzel, E.M. et al. Concerted ESCRT and clathrin recruitment waves define the timing and morphology of intraluminal vesicle formation. Nat Commun 9, 2932 (2018).

29. Luzio, J.P., Pryor, P.R. & Bright, N.A. Lysosomes: fusion and function. Nat Rev Mol Cell Biol 8, 622–632 (2007).

30. Bridgewater, R.E., Norman, J.C. & Caswell, P.T. Integrin trafficking at a glance. Journal of cell science 125, 3695–3701 (2012).

31. Sigismund, S., Avanzato, D. & Lanzetti, L. Emerging functions of the EGFR in cancer. Mol Oncol (2017).

32. Tod, J. et al. Pro-migratory and TGF-beta-activating functions of alphavbeta6 integrin in pancreatic cancer are differentially regulated via an Eps8-dependent GTPase switch. J Pathol (2017).

33. Dong, X. et al. Force interacts with macromolecular structure in activation of TGF-beta. Nature 542, 55–59 (2017).

34. Worthington, J.J., Klementowicz, J.E. & Travis, M.A. TGFß: a sleeping giant awoken by integrins. Trends in biochemical sciences 36, 47–54 (2011).

35. Badouel, C. & McNeill, H. SnapShot: The hippo signaling pathway. Cell 145, 484–484 e481 (2011).

36. Piccolo, S., Dupont, S. & Cordenonsi, M. The biology of YAP/TAZ: hippo signaling and beyond. Physiol Rev 94, 1287–1312 (2014).

37. Elosegui-Artola, A. et al. Force Triggers YAP Nuclear Entry by Regulating Transport across Nuclear Pores. Cell 171, 1397–1410 e1314 (2017).

38. Oria, R. et al. Force loading explains spatial sensing of ligands by cells. Nature 552, 219–224 (2017).

39. Sheppard, D. Integrin-mediated activation of latent transforming growth factor beta. Cancer Metastasis Rev 24, 395–402 (2005).

40. Zanconato, F., Cordenonsi, M. & Piccolo, S. YAP/TAZ at the Roots of Cancer. Cancer Cell 29, 783–803 (2016).

41. Dupont, S. Role of YAP/TAZ in cell-matrix adhesion-mediated signalling and mechanotransduction. Exp Cell Res 343, 42–53 (2016).

42. DuFort, C.C., Paszek, M.J. & Weaver, V.M. Balancing forces: architectural control of mechanotransduction. Nat Rev Mol Cell Biol 12, 308–319 (2011).

43. Levental, K.R. et al. Matrix crosslinking forces tumor progression by enhancing integrin signaling. Cell 139, 891–906 (2009).

44. Diaz-Martin, J. et al. Nuclear TAZ expression associates with the triple-negative phenotype in breast cancer. Endocr Relat Cancer 22, 443–454 (2015).

45. Cordenonsi, M. et al. The Hippo Transducer TAZ Confers Cancer Stem Cell-Related Traits on Breast Cancer Cells. CELL 147, 759–772 (2011).

46. Horton, E.R. et al. Definition of a consensus integrin adhesome and its dynamics during adhesion complex assembly and disassembly. Nat Cell Biol 17, 1577–1587 (2015).

47. Horton, E.R. et al. Modulation of FAK and Src adhesion signaling occurs independently of adhesion complex composition. J Cell Biol 212, 349–364 (2016).

48. Robertson, J. et al. Defining the phospho-adhesome: phosphoproteomic analysis of integrin signalling. Nature Communications, In press (2015).

49. Byron, A. et al. A proteomic approach reveals integrin activation state-dependent control of microtubule cortical targeting. Nat Commun 6, 6135 (2015).

50. Schiller, H.B., Friedel, C.C., Boulegue, C. & Fassler, R. Quantitative proteomics of the integrin adhesome show a myosin II-dependent recruitment of LIM domain proteins. EMBO reports 12, 259–266 (2011).

51. Schiller, H.B. et al. beta1- and alphav-class integrins cooperate to regulate myosin II during rigidity sensing of fibronectin-based microenvironments. Nat Cell Biol 15, 625–636 (2013).

52. Kuo, J.C., Han, X., Hsiao, C.T., Yates, J.R., 3rd & Waterman, C.M. Analysis of the myosin-II-responsive focal adhesion proteome reveals a role for beta-Pix in negative regulation of focal adhesion maturation. Nature cell biology 13, 383–393 (2011).

53. Gutteridge, E. et al. The effects of gefitinib in tamoxifenresistant and hormone-insensitive breast cancer: a phase II study. Int J Cancer 126, 1806–1816 (2010).

54. Osborne, C.K. et al. Gefitinib or Placebo in Combination with Tamoxifen in Patients with Hormone Receptor-Positive Metastatic Breast Cancer: A Randomized Phase II Study. Clinical Cancer Research 17, 1147–1159 (2011).

55. McShane, L.M. et al. REporting recommendations for tumour MARKer prognostic studies (REMARK). Eur J Cancer 41, 1690–1696 (2005).

56. Curtis, C. et al. The genomic and transcriptomic architecture of 2,000 breast tumours reveals novel subgroups. Nature 486, 346–352 (2012).

57. Jones, M.C. et al. Isolation of integrin-based adhesion complexes. Curr Protoc Cell Biol 66, 9 8 1–9 8 15 (2015).

58. Wu, J. et al. Integrated network analysis platform for protein-protein interactions. Nat Methods 6, 75–77 (2009).

59. Robertson, J. et al. Defining the phospho-adhesome through the phosphoproteomic analysis of integrin signalling. Nat Commun 6, 6265 (2015).

60. Zaidel-Bar, R., Itzkovitz, S., Ma’ayan, A., Iyengar, R. & Geiger, B. Functional atlas of the integrin adhesome. Nat Cell Biol 9, 858–867 (2007).

61. Hoopmann, M.R., Merrihew, G.E., von Haller, P.D. & MacCoss, M.J. Post analysis data acquisition for the iterative MS/MS sampling of proteomics mixtures. J Proteome Res 8, 1870– 1875 (2009).

62. Cutillas, P.R. & Vanhaesebroeck, B. Quantitative profile of five murine core proteomes using label-free functional proteomics. Mol Cell Proteomics 6, 1560–1573 (2007).

63. Smyth, G.K. Linear models and empirical bayes methods for assessing differential expression in microarray experiments. Stat Appl Genet Mol Biol 3, Article3 (2004).

64. Eccles, R.L. et al. Bimodal antagonism of PKA signalling by ARHGAP36. Nat Commun 7, 12963 (2016).

65. Rainero E., v.d.B.P.V.E., Norman J.C. Internalisation, Endosomal Trafficking and Recycling of Integrins During Cell Migration and Cancer Invasion, in Vesicle Trafficking in Cancer, Vol. I. (eds. Y. Yarden & G. Tarcic) 327 – 359 (Springer, New York; 2013).

66. Martiel, J.L. et al. Measurement of cell traction forces with ImageJ. Methods Cell Biol 125, 269–287 (2015).

